# Macrophage and neutrophil heterogeneity at single-cell spatial resolution in inflammatory bowel disease

**DOI:** 10.1101/2022.11.28.518139

**Authors:** Alba Garrido-Trigo, Ana M. Corraliza, Marisol Veny, Isabella Dotti, Elisa Melon-Ardanaz, Aina Rill, Helena L. Crowell, Ángel Corbí, Victoria Gudiño, Miriam Esteller, Iris Álvarez-Teubel, Daniel Aguilar, M Carme Masamunt, Emily Killingbeck, Youngmi Kim, Michael Leon, Sudha Visvanathan, Domenica Marchese, Ginevra Caratù, Albert Martin-Cardona, Maria Esteve, Julian Panés, Elena Ricart, Elisabetta Mereu, Holger Heyn, Azucena Salas

**Author notes:** Contributed equally (Elisabetta Mereu, Holger Heyn).

## Abstract

Ulcerative colitis (UC) and Crohn’s disease (CD) are chronic inflammatory intestinal diseases that show a perplexing heterogeneity in manifestations and response to treatment. The molecular basis for this heterogeneity remains uncharacterized. We applied single-cell RNA sequencing and CosMx™ Spatial Molecular Imaging to human colon and found the highest diversity in cellular composition in the myeloid compartment of UC and CD patients. Besides resident macrophage subsets (M0 and M2), patients showed a variety of activated macrophages including classical (M1 CXCL5 and M1 ACOD1) and new inflammation-dependent alternative (IDA) macrophages. In addition, we captured intestinal neutrophils in three transcriptional states. Subepithelial IDA macrophages expressed *NRG1*, which promotes epithelial differentiation. In contrast, *NRG1*^low^ IDA macrophages were expanded within the submucosa and in granulomas, in proximity to abundant inflammatory fibroblasts, which we suggest may promote macrophage activation. We conclude that macrophages sense and respond to unique tissue microenvironments, potentially contributing to patient-to-patient heterogeneity.

## INTRODUCTION

Inflammatory bowel diseases (IBDs) are chronic immune-mediated diseases of the gastrointestinal tract that are normally classified as Crohn’s disease (CD) or ulcerative colitis (UC) based on histological, imaging, and clinical features^1^. Despite this classification, a remarkable degree of variability is routinely observed in clinics in terms of disease severity, response to therapy and disease progression^2^. However, no validated clinical or biological features have been established to explain and faithfully predict such variability.

Single-cell RNA sequencing (scRNA-seq) of the intestinal mucosa recently provided a description of close to sixty different cell types present in UC^3–6^ and ileal CD^7^, emphasizing the magnitude of changes across cell populations in the context of intestinal inflammation. We applied scRNA-seq to colonic biopsies of healthy and active UC and colonic CD patients with a focus on understanding the heterogeneity among patients within each cellular compartment. The myeloid compartment, including macrophages and neutrophils, showed the highest diversity in composition within patient groups, suggesting these cell types may contribute to differences among patients in disease phenotype and progression over time.

Macrophages are resident immune cells that act as gatekeepers in tissues and are well-known for their ability to sense and adapt to environmental changes. Two states of activation were initially described in mice, termed classical or M1 and alternative or M2 macrophages^8^ that express different markers^9–11^. Importantly, signature genes for both subsets have been found in the human colon, including IBD samples^12^, which showed a marked increase in M1 populations in those patients. Macrophage polarization has been more recently revisited showing a broader functional repertoire of this cell population, and important differences among intestinal macrophages phenotype and function have been linked to their spatial distribution within intestinal layers^14^.

By combining scRNA-seq with the recently developed highly multiplexed CosMx Spatially Molecular Imaging (SMI) (NanoString Technologies) analysis^13^, we discovered the signatures and tissue distribution of previously uncharacterized intestinal macrophage populations, including two subsets of resident and novel inflammation-related macrophages. This approach helped us understand the potential *in vivo* roles and likely interacting partners, including epithelial cells and fibroblasts, of the different macrophage subsets. Overall, our study emphasizes the diversity and plasticity of the intestinal myeloid compartment, specifically of the macrophage and neutrophil populations, and reveals novel mechanisms potentially contributing to heterogeneity in IBD.

## RESULTS

### Integration of single-cell RNA sequencing and spatial molecular imaging analysis provides a map of healthy and inflamed colon

ScRNA-seq analysis of colonic biopsies from HC (n=6), CD (n=6) and UC (n=6) active patients (Fig. 1a; Supplementary Table 1) totaling 46,700 cells identified 79 different clusters (Fig. 1b), whose proportions varied significantly between disease groups (Fig. 1b and 1c; Extended data Fig 1a). Each compartment (epithelium, stroma, B and plasma cells, T cells and myeloid cells) was isolated *in silico* to achieve higher resolution on cell populations. Analysis of differentially expressed genes (DEGs) detected cluster-specific marker genes with adjusted p-values (see online Methods). DEG for each cluster (Supplementary Table 2) were used to annotate subpopulations. Subsets, such as inflammatory fibroblasts, neutrophils, or inflammatory M1 macrophages, were found in some CD or UC patients, but were absent in HC (Extended data Fig 1a).

**Figure 1.**
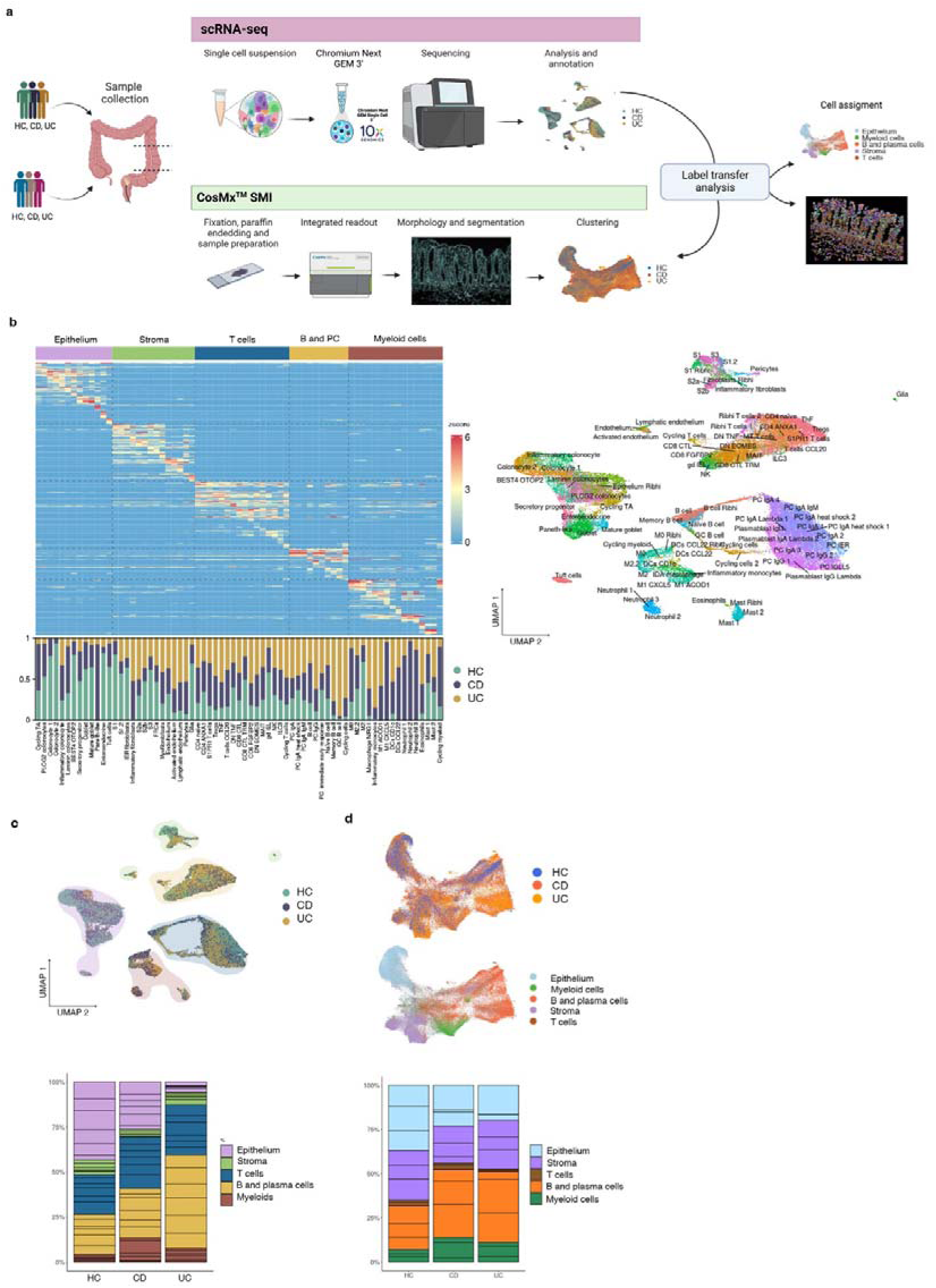
Integration of single cell RNA sequencing (scRNA-seq) and Spatial Molecular Imaging (SMI) provides a map of healthy and inflammatory bowel disease (IBD) colonic biopsies. **a,** Overview of the study design for scRNA-seq, CosMx™ SMI and label transfer from scRNA-seq annotations to the SMI dataset. Two cohorts of Colonic samples including active Crohn’s disease (CD), active ulcerative colitis (UC) and healthy controls (HC) were processed by scRNA-seq (n=18 samples) and CosMx™ SMI (n=9 samples). See Supplementary Table 1 for details. **b,** Heatmap of top marker genes discriminating the different cell subsets (epithelium, stroma, T cells, B and Plasma cells, and myeloid cells) and, below, barplots representing the proportions of each cell type resolved by scRNA-seq for HC, CD and UC. On the right, UMAP showing annotation of all cell types identified by scRNA-seq. **c,** UMAP and barplots of scRNA-seq data. Cells in UMAP are colored by group origin (HC, CD and UC) and clusters are shaded by cell subset (epithelium, stroma, T cells, B and Plasma cells, and myeloid cells). Barplots show the proportions of each cell subset in HC, CD and UC. **d,** UMAP and barplots of CosMx™ SMI data. Top UMAP shows cells colored by group (HC, CD and UC) while bottom UMAP is colored by cell subset (epithelium, stroma, T cells, B and Plasma cells and myeloid cells). Barplots show the proportions of each cell subset in HC, CD and UC.

An additional cohort of formalin-fixed paraffin-embedded (FFPE) colonic samples (Fig. 1a; Supplementary Table 1, cohort 2) was processed using CosMx Spatial Molecular Imaging (SMI; NanoString Technologies)^15^. Scanner for Fields of View (FoVs) and immunofluorescence staining of pan-cell markers (CD45, PanCK, CD3) were performed on all tissues (Extended data Fig 2a and 2b). Selected FoVs were processed by CosMx SMI using a multiplex panel of 1,000 genes. Annotation of cells was performed by label transfer based on scRNA-seq clusters, using the 100 top-ranked markers and count matrix (see online Methods), which assigned a unique label to each cell (Fig. 1d; Extended data Fig 1b and Extended data Fig 2).

Thus, by integrating scRNA-seq and CosMx SMI from human colonic samples, we have generated the first spatially resolved map of healthy and diseased colon at single-cell spatial resolution.

### Transcriptional analysis at single-cell and spatial resolution reveals novel populations of resident and inflammatory macrophages in the colonic mucosa

Remarkably, when comparing cluster proportions within patient groups, the largest discrepancies between individuals were found within the myeloid, followed by the stromal, compartment (Extended data Fig. 1a and 3a), suggesting that the composition of these cell groups may be heavily influenced by patient-dependent factors, and could thus contribute to patient-to-patient heterogeneity.

Pooled HC, CD and UC scRNA-seq data identified different myeloid clusters including several macrophage subtypes, dendritic cells, inflammatory monocytes, mast cells, neutrophils, and eosinophils, whose proportions changed in both UC and CD samples compared to controls (Fig. 2a and Extended data Fig. 3b; Supplementary Table 2). To perform a differential abundance test without relying on cell clustering, we applied Milo a tool which relies on k-nearest neighbor graphs^16^, and we confirmed the differential abundance between groups in the myeloid compartment (Fig. 2b).

**Figure 2.**
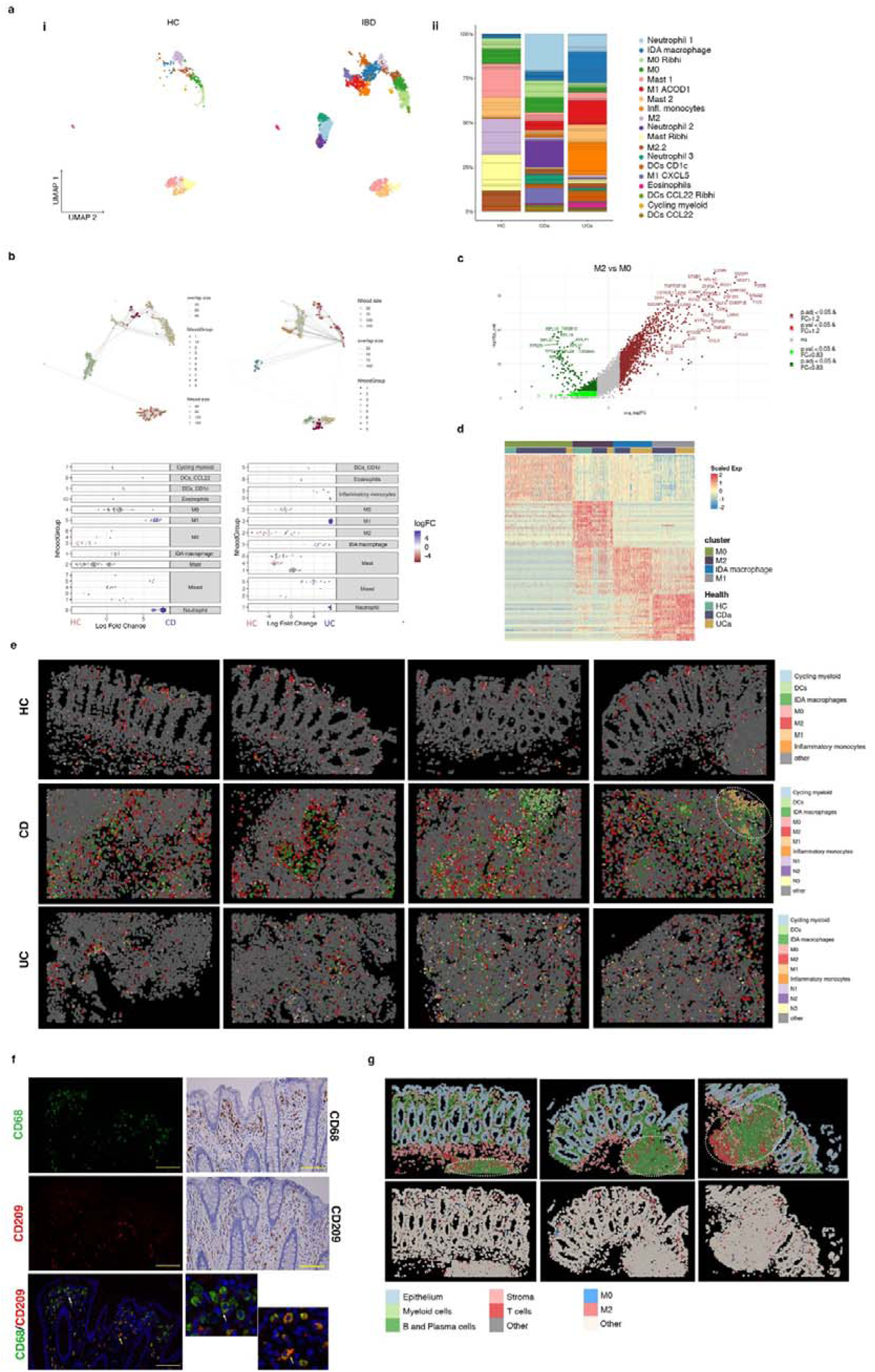
Analysis of myeloid cell subsets in healthy and inflamed colonic mucosa. **a**, UMAP representation of scRNA-seq data showing the different myeloid clusters (macrophages, mast cells, dendritic cells, neutrophils and eosinophils) present in healthy controls (HC, n=6) and IBD colonic samples (CD n= 6 and UC n=6) (i); myeloid cell subset proportions across healthy and IBD samples, x-axis represents patient groups and y-axis the percentages of each myeloid subset (ii). **b**, Cell type enrichment analysis using the differential abundance test Milo. Top plots represent independent clustering performed by Milo, in which nhood groups are shown for HC and CD cells (left) and for HC and UC cells (right). Lower panels show the cell subsets enriched in HC *vs* CD (left) or in HC *vs* UC (right). Annotation of each nhood group based on our analysis (panel a) is shown on the right for each comparison. The logarithm of the fold-change comparing CD or UC to HC is represented on the horizontal axis. The nhood group number is represented in the vertical axis for each analysis. **c**, Volcano plot of differentially expressed genes (DEGs) in M2 compared to M0 macrophages (Supplementary Table 3 contains the complete list of genes). Genes upregulated in M2 macrophages are shown in dark (UUP, p value<0.05) or light red (UP, nominal p value<0.05). Genes downregulated in M2 are shown in dark (DDW, FDR<0.05) or light green (DW, p value<0.05). **d**, Heat map showing the average expression of DEGs for M0, M2, M1 and IDA macrophages in HC, CD and UC. **e**, CosMx™ SMI images showing spatial distribution of the different myeloid cell populations in representative Fields of View (FoVs) of colonic tissue of two HC, one inflamed CD and two inflamed UC patients. White dotted circle indicates ulcerated area with abundant M1 macrophages. **f**, On the left and right insets, double immunofluorescence showing M2 (CD209^+^ in red and CD68^+^ in green) and M0 (CD209^-^ and CD68^+^) subsets in a representative healthy colonic tissue. White and yellow arrows indicate M0 and M2 macrophages, respectively. Right two top panels: immunohistochemistry showing the distribution of CD68 and CD209 markers in healthy colonic tissue (Scale bar 20 µm). **g**, CosMx™ SMI images of a representative healthy colonic mucosa with lymphoid follicles (highlighted by dotted circles). Upper panels show the cellular localization of the five cell subsets (epithelial, stroma, T cells B and plasma cells, and myeloid cells) and lower panels M0 and M2 resident macrophages location.

In healthy colon, resident macrophages (expressing *C1QA*, *HLA-DRB1* and *SELENOP*, among others) were found in different transcriptional states (Extended data Fig. 3b). These included macrophages expressing well-described M2-specific markers (i.e., *CD163L1*, *CD209*, *FOLR2*), annotated as M2 and M2.2, and hereafter referred to as M2. We annotated the other two clusters present in HC as M0 (M0 and M0_Rib^hi^), as they lacked M2 markers but highly expressed all other macrophage-specific genes (Fig. 2c and Supplementary Table 3), while showing low expression of *PTPRC* (CD45) (Extended data Fig. 3c). Besides M0 and M2, samples from UC and CD contained inflammatory monocytes and activated macrophage clusters that we annotated as M1 ACOD1, M1 CXCL5 (pooled as M1) and Inflammation-Dependent Alternative (IDA) macrophages (Fig. 2a and 2d).

All myeloid subsets were found in tissue sections analyzed by CosMx SMI, showing marked differences in abundance and spatial distribution depending on the patient and/or disease type (Fig. 2e).

### M0 and M2 represent two independent states

Until now, M0 macrophages have not been formally described in the human intestine. Thus, we first compared our monocyte/macrophage signatures to publicly available data from HC, UC ^3^ and CD terminal ileum^7^ (Extended data Fig. 4a) and found, in both cohorts, populations of macrophages that resembled the M0 and M2 subsets (Jaccard indexes=0.3) (Extended data Fig. 4b).

In agreement with the scRNA-seq data, we could visualize both CD209^+^CD68^+^ (M2) and CD209^-^ CD68^+^ (M0) cells using immunostaining in healthy colonic lamina propria, mostly localizing below the apical epithelium (Fig. 2f) and also present throughout the lamina propria. Likewise, M0 and M2 cells were identified by CosMx SMI analysis (Fig. 2), confirming the dual identity of these resident macrophages.

To understand their phylogenetic origin and their relation to other previously described macrophage subsets, we mapped our dataset to a recently published human monocyte-macrophage database containing data from 41 studies on several organs and diseases (MoMac-VERSE)^17^ (Extended data Fig. 4c). M0, M2.2 and M2 mapped to independent macrophage clusters within the MoMac-Verse dataset, supporting the hypothesis that they do represent two unique states.

Indeed, comparison of M0 and M2 macrophages in our dataset to *in vitro* monocyte-derived macrophages from published datasets^18^ showed high similarity between intestinal M2 and *in vitro* M-CSF monocyte-derived macrophages (Extended Fig. 5a), while no or little overlapping with M0 macrophages was observed under these same conditions.

Interestingly, trajectory analysis of our data (Extended data Fig. 5b) and the two public datasets annotated above (Extended data Fig. 4d) suggests separate pseudo-time states for M0 and M2 clusters. In our data, M2.2 clusters, which express M2 markers, appear close to M0, suggesting they may represent a transitional state between the two resident compartments.

Overall, we conclude that in the healthy colon, resident macrophages are found in at least two states. M2 macrophages could potentially originate from circulating monocytes exposed to M-CSF in tissues, while the origin of M0 macrophages remains unknown.

### Inflammation-associated macrophages, including M1 and novel Inflammation-Dependent Alternative macrophages, show highly heterogeneous signatures among patients

Compared to HC, IBD patients showed a marked increase in the total number and transcriptional heterogeneity of the macrophage population (Fig. 2a and Extended Data Fig. 1). Apart from of M0 and M2, we found inflammatory/activated macrophages in at least three different states: two transcriptionally different M1 populations (M1 ACOD1 and M1 CXCL5) and a newly identified IDA macrophage cluster, in addition to a population of inflammatory monocytes. Comparison of these inflammation-associated cell types to the MoMac VERSE data set^17^ is also shown in Extended data Fig. 4d.

In contrast to M0 and M2 macrophages, the similarities between inflammatory macrophages in our cohort and those found in other intestinal datasets^3,7^ were weaker, suggesting that activated macrophages may be found in highly patient/context-dependent states (Jaccard Index <=0.14 Smillie et al, and Jaccard Index <=0.16 Martin et al.; Extended data Fig. 4c).

As with M2 and M-CSF-derived macrophages, intestinal M1 CXCL5 cells showed high similarity to the *in vitro* GM-CSF-derived macrophages, while the signature of M1 ACOD1 was shared by both M-CSF and GM-CSF-derived macrophages stimulated with LPS^18^ (Extended data Fig. 5c). Remarkably, IDA macrophages showed the most transcriptional similarity to M-CSF-derived macrophages treated with serotonin (5-HT) ^19^(Extended data Fig. 5c).

Based on trajectory analysis, M1 subsets populated a different branch to those of M2/M0 subsets in all 3 datasets (Extended data Fig. 4d and 5b), with inflammatory monocytes exclusively transitioning towards the fully activated M1, state. IDA macrophages instead appear to contain a heterogeneous population divided between the M1 and the M2 branches, suggesting they may represent a transitional state between those subsets. Analysis of overlapping markers between M1, M2 and IDA macrophages reveals that the latter share about 16% and 10% of its top 200 marker genes with M2 and M1, respectively (Extended Data Fig. 5d).

Overall, we show that in the context of inflammation, macrophages can adopt diverse transcriptional signatures, with high heterogeneity between patients. Our data also suggests that intestinal macrophages could originate from monocytes activated under different stimuli including GM-CSF, GM-CSF+LPS, M-CSF+LPS or M-CSF+5-HT, highlighting the importance of the microenvironment in modulating their phenotypes.

### Inflammation-dependent alternative macrophages express neuregulin 1

Markers of IDA macrophages include epidermal growth factor receptor (*EGFR*) family ligands like AREG and *HBEGF* and specifically *NRG1*, while showing lower expression of M1 and M2 canonical markers (Fig. 3a and Supplementary Table 2).

**Figure 3.**
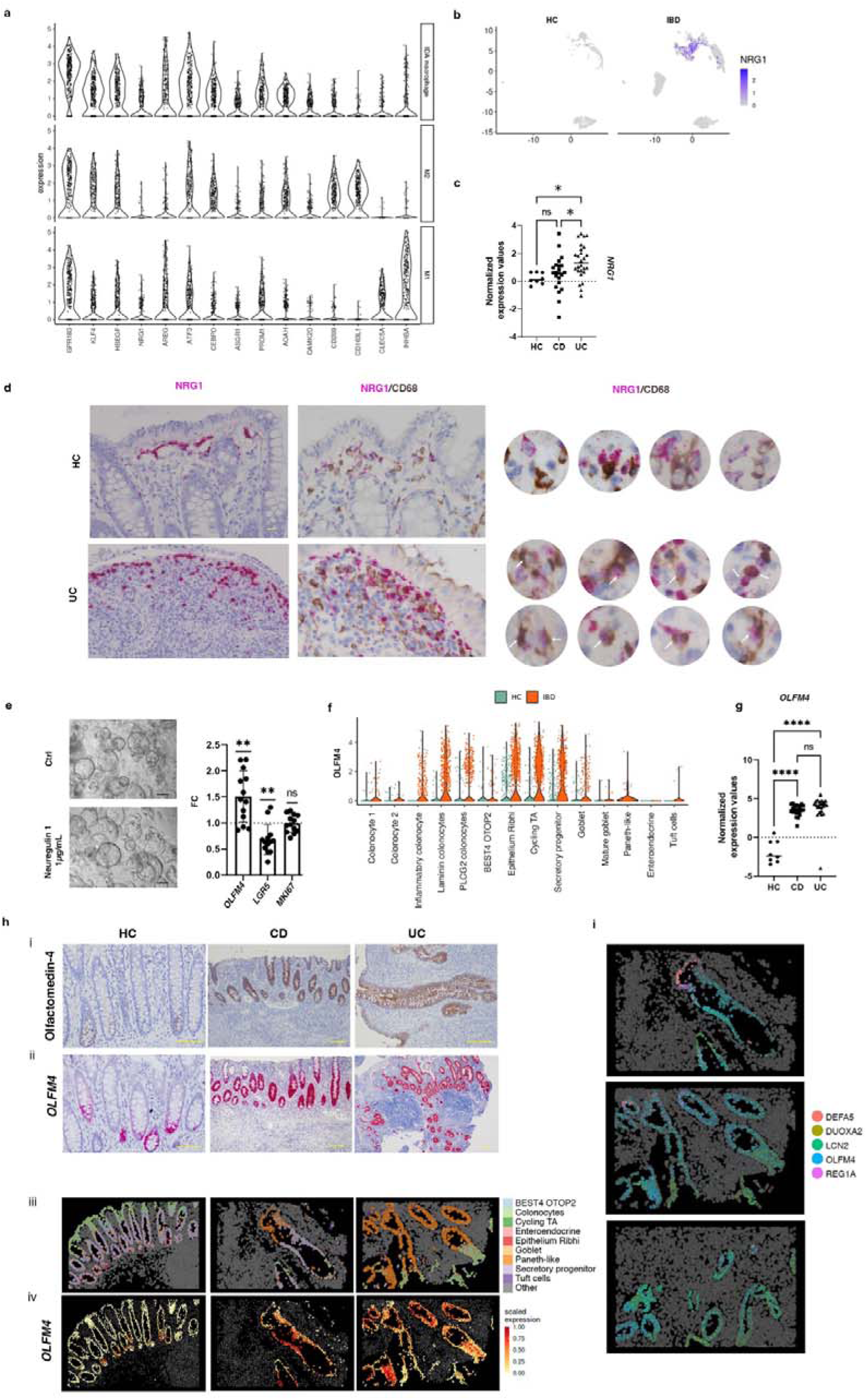
Neuregulin 1 expression and function in colonic mucosa. **a**, Violin plot showing the expression of marker genes of IDA, M2 and M1 macrophages from pooled HC, CD and UC scRNA-seq data. **b**, UMAP showing *NRG1* expression in the myeloid compartment of HC and IBD (CD and UC) data. **c**, *NRG1* expression from bulk biopsy RNA-seq data in HC (n=8), and active CD (n=22) and UC (n=26) patients. Ordinary one-way ANOVA corrected (p<0,05(*), ns>0,01). **d**, Double *In situ* hybridization of *NRG1* and immunohistochemistry for CD68 in HC and active UC tissue. CD68^+^ NRG1^-^, CD68^+^ NRG1^+^ and CD68^-^NRG1^+^ cells are shown. White arrows show NRG1^+^ CD68^+^ cells in UC patients (Scale bar= 10 µm) **e**, Representative pictures of human-derived epithelial organoids under vehicle (Ctrl) or Neuregulin 1 (1 μg/mL) for 48h. Scale bars= 100 µm. mRNA expression of *OLFM4*, *LGR5* and *MKI67* was determined by RT-qPCR (n=13). Data is shown as fold change (FC) relative to the vehicle treated condition. Bars represent mean ± standard deviation (SD). p<0,05(*), p<0,01 (**), p<0,001(***), ns: not significant. **f**, Violin plot showing *OLFM4* expression (y-axis) in all epithelial cell subsets (x-axis) in HC (green) and IBD (orange) samples by scRNA-seq data. **g**, *OLFM4* expression by bulk RNA-seq of biopsy samples from HC (n=8), active CD (n=22) and active UC (n=26) patients. p<0,05(*), p<0,01 (**), p<0,001(***), p<0,0001(****), ns: not significant. **h**, (i) Olfactomedin 4 immunostaining and (ii) *OLFM4 in situ* hybridization in HC, active CD and active UC colon. Scale bars= 100 µm. (iii) CosMx™ SMI visualization and localization of the different epithelial cell subsets described by scRNA-seq, from left to right in a HC and two UC representative Fields of View (FoVs) and (iv) mean expression of *OLFM4* in each of those cells analyzed by CosMx™ SMI **i**, Expression of *DEFA5*, *LCN2*, *DUOX2A*, *REG1A*, and *OLFM4* within epithelium of representative FoVs of a UC patient analyzed by CosMx™ SMI. Dots represents mRNA molecules.

In agreement with scRNA-seq data (Fig. 3b), *NRG1* was significantly increased in bulk RNA-seq analysis of UC colonic mucosa compared to HC and CD (Fig. 3c), suggesting the potential involvement of neuregulin 1 in these patients. *In situ* hybridization of *NRG1* mRNA confirmed its expression on abundant CD68^+^ macrophages in IBD (Fig. 3d). In contrast, *NRG1* expression in HC was more specific to a population underlying the surface epithelium, with no or little colocalization within CD68^+^ macrophages (Fig. 3d). Indeed, our scRNA-seq and CosMx SMI analysis showed that in HC mucosa, S2b (*SOX6*^+^) (localized at the most apical area), but not S2a pericryptal fibroblasts (Extended data Fig. 6d and 6e), also express *NRG1* (Extended Data Fig. 6a, 6b and 6c). In addition, fibroblast-expression of *NRG1* was also markedly increased in UC (Extended data Fig. 6b and 6f).

Neuregulin 1 binds to ErbB receptors^20^ and can promote epithelial differentiation to secretory lineages in human ^21^ and stem cell proliferation and regeneration in mice^22^ Using intestinal epithelial stem cell-enriched organoids, we found that neuregulin 1 significantly decreased expression of the stem cell marker *LRG5* and upregulated *OLFM4*, despite inducing no changes in the proliferation marker *MKI67*(Fig. 3e). Expression of OLFM4 is mostly restricted to a progenitor cell type in healthy tissues (Epithelium Ribhi, Secretory Progenitors) (Fig. 3f; Extended Data Fig. 7a, 7b and 7c), while it is dramatically increased in UC and CD, as shown by scRNAseq (Fig. 3f), bulk RNA analysis (Fig. 3g), immunostaining, *in situ* hybridization, and CosMx™ SMI of colonic tissue (Fig. 3h), in agreement with previous data^23^. Epithelial cell populations by CosMx SMI in a HC Field of View (FoV) are shown in Extended Data Fig.7c

Besides *OLFM4*, through SMI we observed other changes that occur in the intestinal epithelium of IBD patients, including the upregulation of anti-microbial mechanisms such as the expression of defensins (*DEFA5*), lipocalins (*LCN2*) and enzymes involved in producing reactive oxygen species (*DUOXA2*) (Fig. 3i; Extended Data Fig. 7d and 7e).

In summary, we show that IDA macrophages and S2b fibroblasts overexpress *NRG1* in IBD, particularly UC patients. Neuregulin 1, among other roles, promotes the expansion of the transit-amplifying epithelial compartment, which could play a role in the regeneration of the epithelium.

### CosMx Spatial Molecular Imaging analysis confirms the expansion of Inflammation-Dependent Alternative macrophages and reveals their tissue distribution in inflammatory bowel disease colon

CosMx SMI analysis localized abundant IDA macrophages scattered throughout the inflamed (UC and CD) colon, representing the most expanded inflammation-dependent macrophage state (Fig. 4a), while M1 macrophages were less abundant in the lamina propria and submucosa, but predominated within surface ulcers (Fig. 2g). Of note, in one CD patient we found abundant granulomas (Fig. 4b and Extended Data Fig. 8a), which are aggregates of macrophages, including multiploidy macrophages, which develop in response to persistent inflammation and that are a pathological feature found in about one-fourth of CD patients^24^. IDA macrophages, together with some M2, and a few M0 and M1 macrophages, were the predominant macrophage state within granulomas (Fig. 4b), which were surrounded by diverse lymphoid subsets (Extended data Fig. 8b). The cellular composition of a non-granuloma lymphoid aggregate in the same patient is shown for comparison (Extended data Fig. 8c).

**Figure 4.**
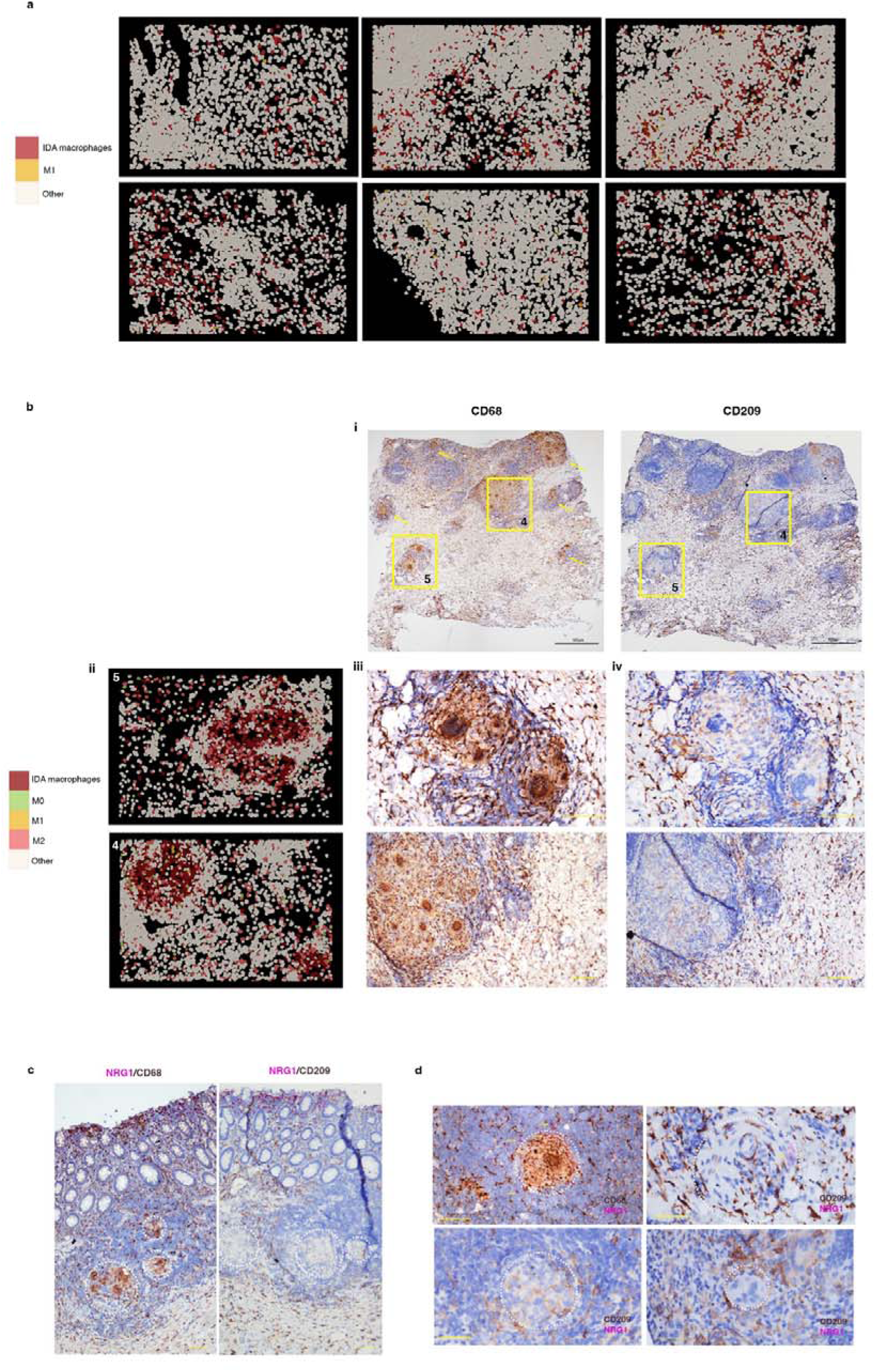
Inflammation-Dependent Alternative (IDA) macrophages are widely distributed in ulcerative colitis (UC) and present in Crohn’s disease (CD) granulomas. **a,** CosMx™ SMI distribution of IDA and M1 macrophages in IBD colonic samples. **b,** CD colonic sample with multiple granulomas (CD b patient). **(i)** Field of Views (FoVs) 4 and 5 from this tissue section are indicated by squares and other granulomas found in the same sample by yellow arrows. **(ii)** Macrophages within granulomas are shown by CosMx™ SMI in FoVs 4 and 5 and protein expression of **(iii)** CD68 and (iv) CD209 is shown by immunohistochemistry on the same tissue sections (scale bars= 100 µm). **c,** Double *NRG1 in situ* hybridization and CD68 or CD209 immunostaining in tissue sections from the CD patient (CD b) containing abundant granulomas. *In situ* hybridization of *NRG1* shows an increasing gradient of expression towards the apical mucosa. Granulomas are indicated by dotted circles (scale bars 100 µm). **d,** Magnified pictures of representative granulomas of the same CD tissue stained for *NRG1* using *in situ* hybridization and CD68 or CD209 immunostaining. Granulomas are indicated by dotted circles and *NRG1* positive cells are shown by arrows (scale bars 100 µm).

In agreement with the CosMx SMI results, immunostaining showed low and scattered staining of the M2 (CD209) markers within CD68^+^ cells in granuloma (Fig. 4b, 4c and Extended data Fig.8a). Compared to lamina propria macrophages, *NRG1* expression within the granulomas was low (Fig. 4c).

Thus, IDA macrophages abundantly present in the inflamed colon display differential NRG1 expression depending on their tissue location. While *NRG1*^hi^ IDA macrophages localize to the most apical subepithelial compartment of the mucosa, *NRG1*^low^ alternatively activated macrophages accumulate within granulomas in CD and in the submucosa of both UC and CD patients, suggesting independent functions.

### Inflammatory fibroblasts co-localize with Inflammation-Dependent Alternative macrophages in inflammatory bowel disease

Given the abundant number and heterogeneity in distribution patterns of IDA macrophages in UC and CD patients, we leveraged the multiplexed spatial data to identify the cell types that were most frequently found in their proximity. IDA macrophages tended to localize near to other macrophage subsets (M0, M2 and M1), some stromal cells, and T cells, particularly CD8^+^ T cells, Tregs and T cells CCL20 (Fig. 5a). Within the stromal compartment, IDA macrophages presented high spatial correlation with inflammatory fibroblasts in both UC and CD (Fig. 5b and 5c), including within granulomas (Extended data Fig 9). Inflammatory fibroblasts were described in UC^5^ and found here in colonic CD (Fig. 5d), as confirmed by biopsy bulk RNAseq (Fig. 5e). Importantly, inflammatory fibroblasts expressed *CSF2* and *CSF3*, encoding for GM-CSF and G-CSF, respectively, and prostaglandin-producing enzymes *PTGS1*, *PTGES*, *PTGS2* (Fig 5f). In fact, a recent study^25^ showed that prostaglandins are produced by activated fibroblasts and drive the differentiation of IDA-like macrophages expressing *HBEGF* and *EREG*, but not *NRG1*, in the synovium of rheumatoid arthritis patients.

**Figure 5.**
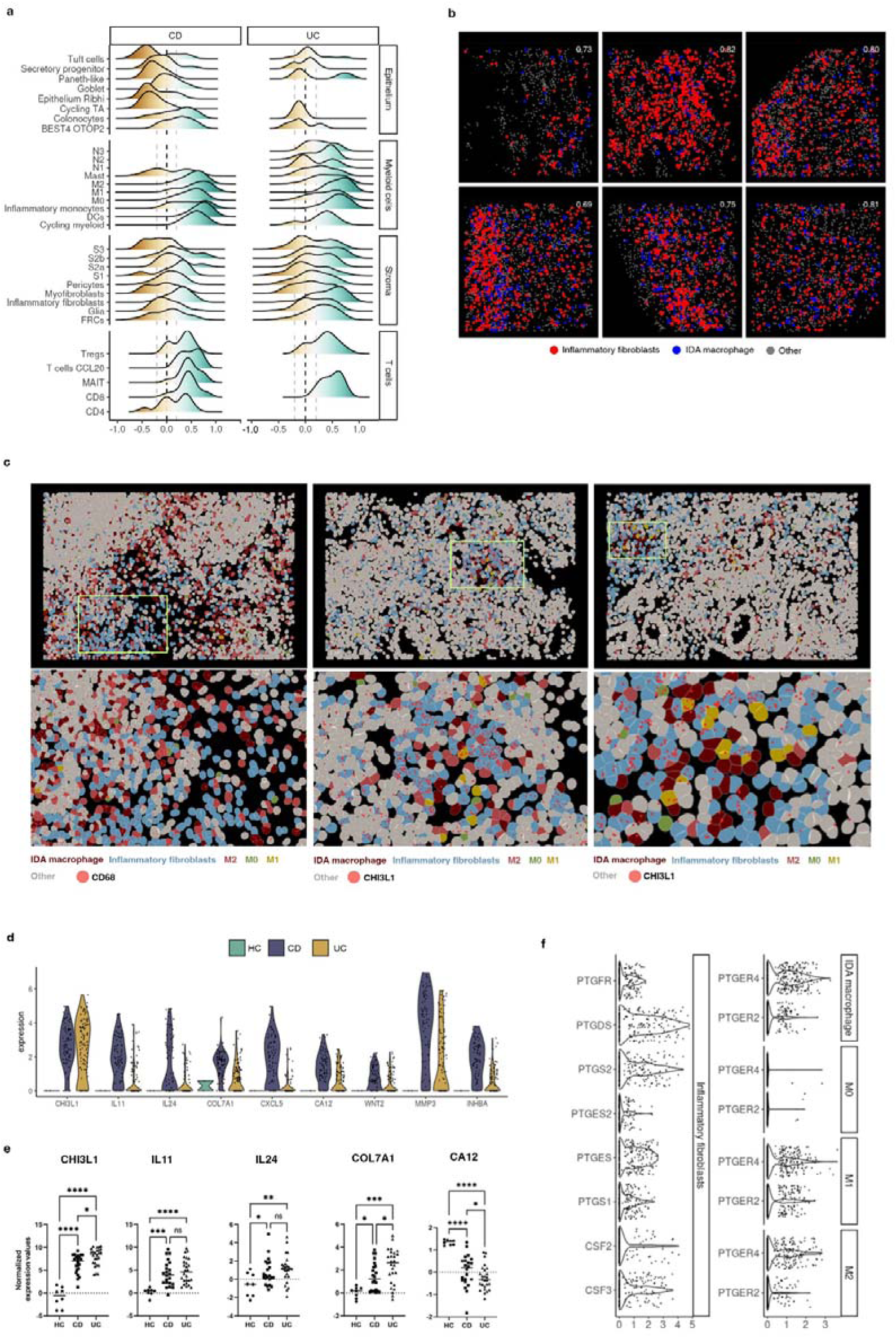
Inflammation-Dependent Associated (IDA) macrophages co-localize with inflammatory fibroblasts. **a**, Ridge plot of co-localization analysis of IDA macrophages and epithelial, other myeloid, stromal and T lymphocytes by CosMx™ SMI. Correlation for cell positions was calculated per cell type (0 indicates no correlation, >1 indicates co-localization with 1 being cells sharing the same position; <1 indicates negative correlation between the indicated cell types). **b**, Representative Fields of View (FoVs) of co-localization analysis between IDA macrophages and inflammatory fibroblasts in inflamed UC tissue. Co-localization scores are indicated in white for each FoV. **c**, Representative FoVs of IBD inflamed tissues containing IDA macrophages and inflammatory fibroblasts. Expression of *CD68* (macrophages) or *CHI3L1* (inflammatory fibroblasts) is shown as red dots. Each dot represents a single mRNA molecule. **d**, Violin plots showing expression (y-axis) of marker genes (x-axis) of inflammatory fibroblasts in HC, active CD and active UC determined by scRNA-seq. **e**, Expression of markers of inflammatory fibroblasts in HC (n=8), and active CD (n=22) and UC (n=26) patients in bulk biopsy RNA-seq data. p<0,05(*), p<0,01 (**), p<0,001(***), p<0,0001(****), ns: not significant. **f**, Violin plot visualizing scRNAseq-based expression (y-axis) of prostaglandin-related genes in inflammatory fibroblasts, IDA macrophage, M2 (M2 & M2.2) and M1 (M1 ACOD1 & M1 CXCL5) in pooled data of HC, CD and UC. Expression of CSF3 and CSF2 in inflammatory fibroblasts has been also included.

Thus, we argue that a crosstalk between inflammatory fibroblasts and macrophages may take place in IBD via specific ligand-receptor interactions.

### Single-cell RNA sequencing reveals a marked heterogeneity of tissue neutrophils in inflammatory bowel disease colonic mucosa

Finally, in addition to the heterogeneity within macrophages in IBD, we also found diverse populations of intestinal granulocytes using scRNA-seq (Fig.2a). Granulocytes, including eosinophils and neutrophils, increased in IBD (Extended Data Fig 10a, 10b and 10c) and expressed distinct membrane protein markers (*CD62L*, *CD193*, *CD69*) compared to their peripheral counterparts, indicating different states of activation (Extended Data Fig 10d). Specific eosinophil markers included *CLC*, *MS4A3*, *CCR3* and the “Th2” cytokines *IL4* and *IL13* (Supplementary Table 2, Extended Data Fig. 3a), while the *HCAR3*, *FCGR3B* (CD16b), *CMTM2* and *PROK2* were specific neutrophil markers (Fig. 2b). Supporting their increase in UC and CD, the expression of most eosinophil and neutrophil markers was significantly increased in bulk biopsy RNAseq (Extended data Fig 10e).

Intestinal neutrophils were found in 3 unique states (annotated as N1, N2 and N3) whose relative abundance varied on individual patients and disease type (Fig. 6a). Compared to N1 and N3, N2 neutrophils, instead expressed higher levels of *CCL3*, *LGALS3* and *CXCR4* (Fig. 6b and c), while N3 neutrophils displayed a marked IFN-response signature (e.g. *GBP1, IRF1* and *FCGR1A)*).

**Figure 6.**
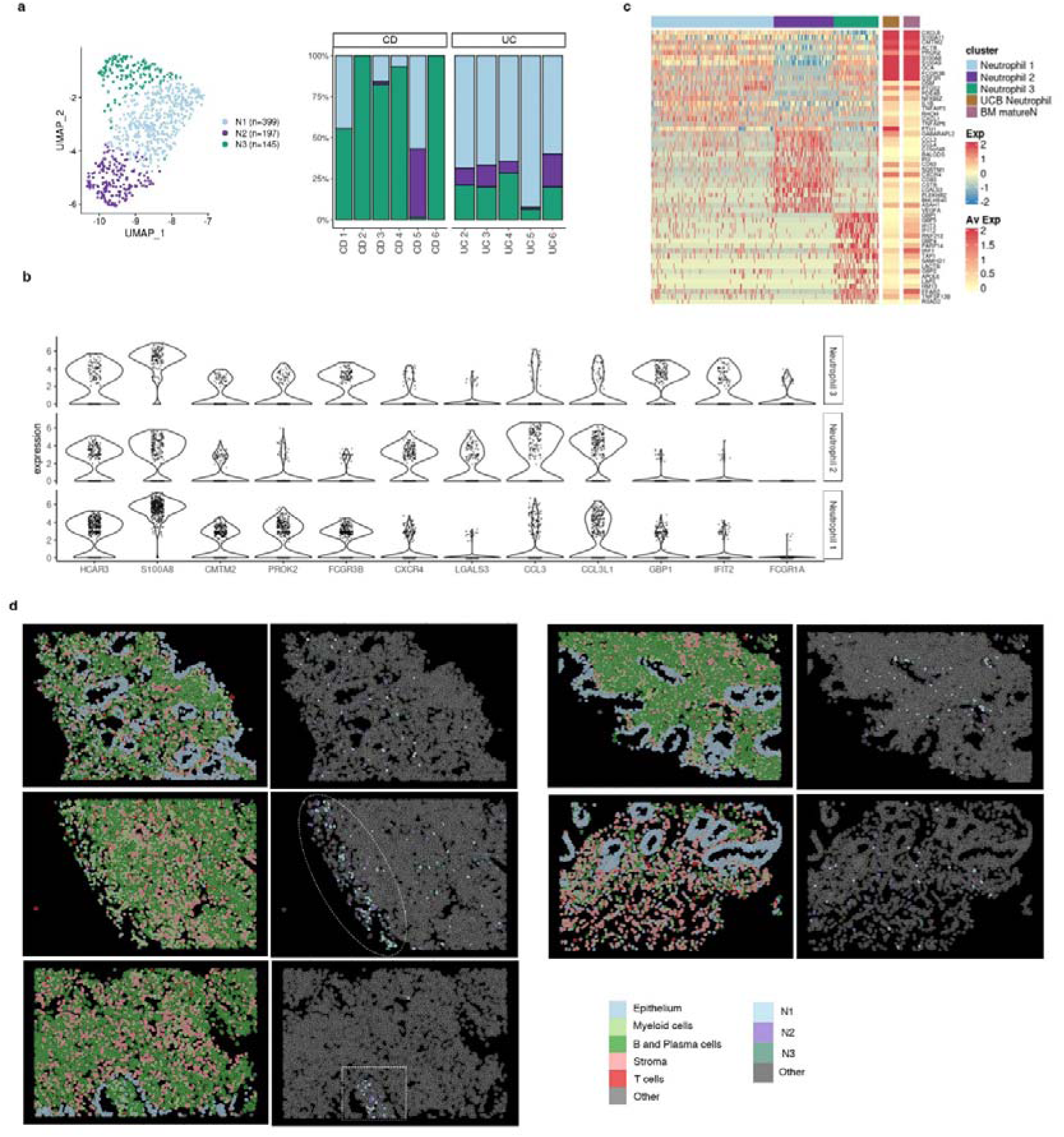
Analysis of the heterogeneity of neutrophil populations in inflammatory bowel disease (IBD) colonic mucosa. **a**, UMAP showing the three neutrophil subsets/states (N1, N2, N3) observed in IBD samples by scRNA-seq analysis. Barplot representing the proportions of each neutrophil subset across health and IBD. **b**, Violin plots visualizing the expression (x-axis) of marker genes common and specific for all three neutrophil populations (y-axis). **c**, Heat map showing the average normalized and scaled expression of differentially expressed genes in all three neutrophil subsets. Average expression of these genes on neutrophils from cord blood and bone marrow mature neutrophils is shown on the far right (Xie, X.*, et al* 2021, Zhao, Y.*, et al* 2019). **d**, Representative CosMx SMI images of IBD inflamed tissue showing the spatial location of N1, N2 and N3 neutrophil subsets. Circle shows the surface of an ulcer, and a square shape is used to indicate a crypt abscess.

Compared to public scRNA-seq datasets from peripheral neutrophils, N1 and N3 neutrophils showed the highest similarity to both bone marrow mature neutrophils (BM matureN)^26^ and to umbilical cord blood neutrophils (UCB)^27^ (Fig. 6c). In contrast, N2 neutrophils showed little overlap with peripheral neutrophils and instead expressed genes suggestive of different tissular locations (e.g., chemokines and receptors, as mentioned above) and different activation states/functions (*CD83*, *CD63*, *FTH1*, *VEGFA*). Protein expression of two of these N2 markers (CXCR4 and CD63) was also confirmed for 10% and 61% of tissue neutrophils compared to 0.26% and 14% of blood neutrophils, respectively (Extended Data Fig. 10d).

All 3 neutrophil subsets were found using CosMx SMI and showed scattered distribution throughout inflamed lamina propria, with predominant localization in crypt abscesses and ulcerated areas (Fig. 6d).

## DISCUSSION

ScRNA-seq has boosted the resolution at which complex tissues, including the inflamed intestine, can be studied^3–7^. Nonetheless, available scRNA-seq datasets lack information on tissue distribution and spatially relevant cell-to-cell interactions. To fill this critical gap, highly multiplexed spatial technologies are rapidly evolving^28^. Our study is the first to provide combined scRNA-seq data with spatial transcriptomics at single-cell resolution to start unraveling patient-dependent disease mechanisms.

We focused on the myeloid compartment, including both macrophage and neutrophil subsets, as they showed the highest degree of variation within patient groups. We argued that changes in these populations may explain disease heterogeneity. Macrophages are well-known for their tissue plasticity. The origin, phenotype, and function of intestinal macrophages, however, continues to be a subject of debate^29,30^. Nonetheless, resident macrophages are known to display heterogenous functions^14,31^. In the context of IBD, activated subsets have been described as having a proinflammatory function^12,32^. Cell classification, however, relies on surface markers that may not be consistently used across studies, thus making standardization challenging. ScRNA-seq provides instead unbiased whole transcriptome profiles of cell types, independently of prior knowledge of marker expression. Using unsorted cells, we discovered at least two unique resident macrophage states (M0 and M2) in healthy colonic mucosa. Both subsets were still present in active patients, together with a variety of activated inflammatory macrophages. Remarkably, the profiles of M0 and M2 macrophages were consistently found in two independent datasets^3,7^ and localized by CosMx SMI to the intestinal lamina propria. In contrast, the transcriptional signatures of inflammation-associated macrophages varied markedly between patients and datasets. We argue that, compared to canonically differentiated resident macrophages, inflammatory macrophages adapt their phenotype to a variety of patient-dependent microenvironments. Based on published data from *in vitro* differentiated macrophages and pseudo-time analysis of scRNA-seq signatures, we also propose that both infiltrating monocytes (found in inflamed samples) and resident M2 macrophages may give rise to activated macrophages. Multiplexed spatial analysis confirmed the diversity in the macrophage populations, and importantly showed that most inflammation-dependent macrophages do not display the full characteristic M1-signature exhibiting instead an alternative activation pattern characterized by the expression of EGFR ligands, *NRG1* and *HBEGF*, and the C-type lectin receptors *CLEC10A* and *ASGR1*. While an M1 signature can be reproduced *in vitro* by exposure of blood monocytes to GM-CSF, GM-CSF/LPS or M-CSF/LPS, the origin of IDA macrophages remains incompletely understood. We found that an endogenously produced factor, serotonin, which is highly abundant in the gut, can induce a signature on M-CSF-derived macrophages (M2-like) that resembles that of the IDA subset found in IBD. Previous studies have shown that serotonin, primarily produced by enterochromaffin cells, modulates macrophage cytokine secretion and its phenotype *in vitro* and that M2 macrophages, in contrast to M1 macrophages, express the serotonin receptors *HTR1D*, *HTR2B* and *HTR7*^33^. While release of platelet-stored serotonin, can happen via multiple mechanisms at inflammatory sites, our data does not prove this to be a relevant mechanism in patients, nor does it rule out the existence of other signals, including fibroblast-derived prostaglandins^25^, that could drive this alternative activation. To our knowledge, macrophages expressing neuregulin 1 have only been described in a murine model of myocardial infarction^34^and suggested to prevent the progression of fibrosis in mouse hearts. The functional role of IDA macrophages in our patients, thus, remains unclear. *NRG1^hi^* IDA macrophages tended to localize to the most apical side of the mucosa and could potentially play a role in epithelial regeneration based on their ability to produce EGFR ligands, which can act on the intestinal epithelium and drive transition towards a regenerative (*OLFM4*-expressing) phenotype. In contrast, *NRG1^low^* IDA macrophages were found within granulomas of a CD patients and in the submucosa of inflamed patients, suggesting IDA macrophages may play different roles depending on their environment.

Based on both scRNA-seq and SMI, we hypothesize that the interaction between M2 or IDA macrophages and inflammatory fibroblasts could play a role in disease pathophysiology. While there is abundant data in the literature to support the interaction of macrophages and fibroblasts, particularly in the context of cancer and fibrosis^35^, little is known about their crosstalk in the context of chronic inflammation. Inflammatory fibroblasts represent a disease-specific fibroblast subset characterized by the expression of multiple cytokines including profibrotic IL11, IL24, IL8, IL6, TGFβ1, and tissue remodeling metallopeptidases, making them attractive therapeutic targets to treat inflammation and potentially, fibrosis, a common and difficult-to-treat complication of chronic intestinal inflammation. Emphasizing their interaction with macrophages, inflammatory fibroblasts express *CSF2* (GM-CSF), which promotes macrophage activation, while activated macrophages can produce mediators (i.e., OSM, IL6, TNF, etc) that can drive fibroblast activation. Furthermore, besides its role on epithelial regeneration, EGFR signaling is a robust regulator of fibroblast motility^36^ and may be involved in cartilage and bone destruction in rheumatoid arthritis^25^.

Beyond macrophages, recent scRNA-seq studies have explored the diversity and plasticity of blood neutrophils^37,38^. These short-lived cells, originally thought to exist in fixed states, have more recently been shown to be transcriptionally dynamic, adopting multiple transcriptional states depending on their maturation stage. Ours is the first report to provide scRNA-seq data on intestinal neutrophils. Compared to available data on periphery, intestinal neutrophils, including a subset that shows a signature of IFN-inducible genes, express the maturation marker CXCR2^37^. Remarkably, CXCR4 neutrophils (N2), which also expressed VEGFA (data not shown), were not found in peripheral datasets. CXCR4, which is essential for bone marrow retention of immature neutrophils, has been identified in mice to mark a subset of pro-angiogenic neutrophils found both in lung and intestine^39^, and to be expressed by neutrophils in inflamed tissues^40,41^. While evidence of angiogenic neutrophils in humans remains elusive, our data points towards the presence of this neutrophil subset, at least in inflamed tissues. Further analysis is required to fully understand the origin and function of these neutrophils in the gut.

Despite the important information that can be drawn from our datasets, there are a few limitations to our study that must be considered. First, the number of total individuals/samples analyzed, especially given the high heterogeneity observed, is too low for us to explore the relationship between the identified signatures and disease behavior. Nonetheless, preliminary analysis of additional datasets, including 74 total patients generated in our group, confirms the presence of heterogeneous resident and inflammatory macrophages, as well as neutrophil subsets across patients. In addition, the 1000-gene SMI panel used in our study, while sufficiently large to cover most cell types, lacked important markers that may have limited our accuracy when assigning cell identities. This may be especially true for cell subsets sharing most of their transcriptomic signature (i.e., N1, N2 and N3 neutrophils).

Overall, we provide evidence to support high patient-dependent heterogeneity within the myeloid compartment in both UC and colonic CD. We argue that intestinal macrophages, which sense changes in the microenvironment, could act as reliable indicators of patient-specific molecular patterns and thus, promising targets. Furthermore, we show that by combining scRNA-seq with SMI, cell subsets can be assigned to likely interacting partners, thus providing crucial niche information. This spatial resolution will be essential in understanding cellular function, and to faithfully link biologically relevant interactions to specific cell types.

## ACKNOWLEDGEMENTS

We thank the Pathology Departments at Hospital Clinic Barcelona and the Biobank Facility and Flow Cytometry unit at the Institut d’Investigacions Biomèdiques August Pi i Sunyer (IDIBAPS) for providing us with the samples required to conduct this study and for the technical help, and all patients for their selfless participation. We also thank Agnès Fernandez-Clotet and Maite Rodrigo Calvo for support, and Joe Moore for editorial assistance.

---

**Extended Data Figure 1.**
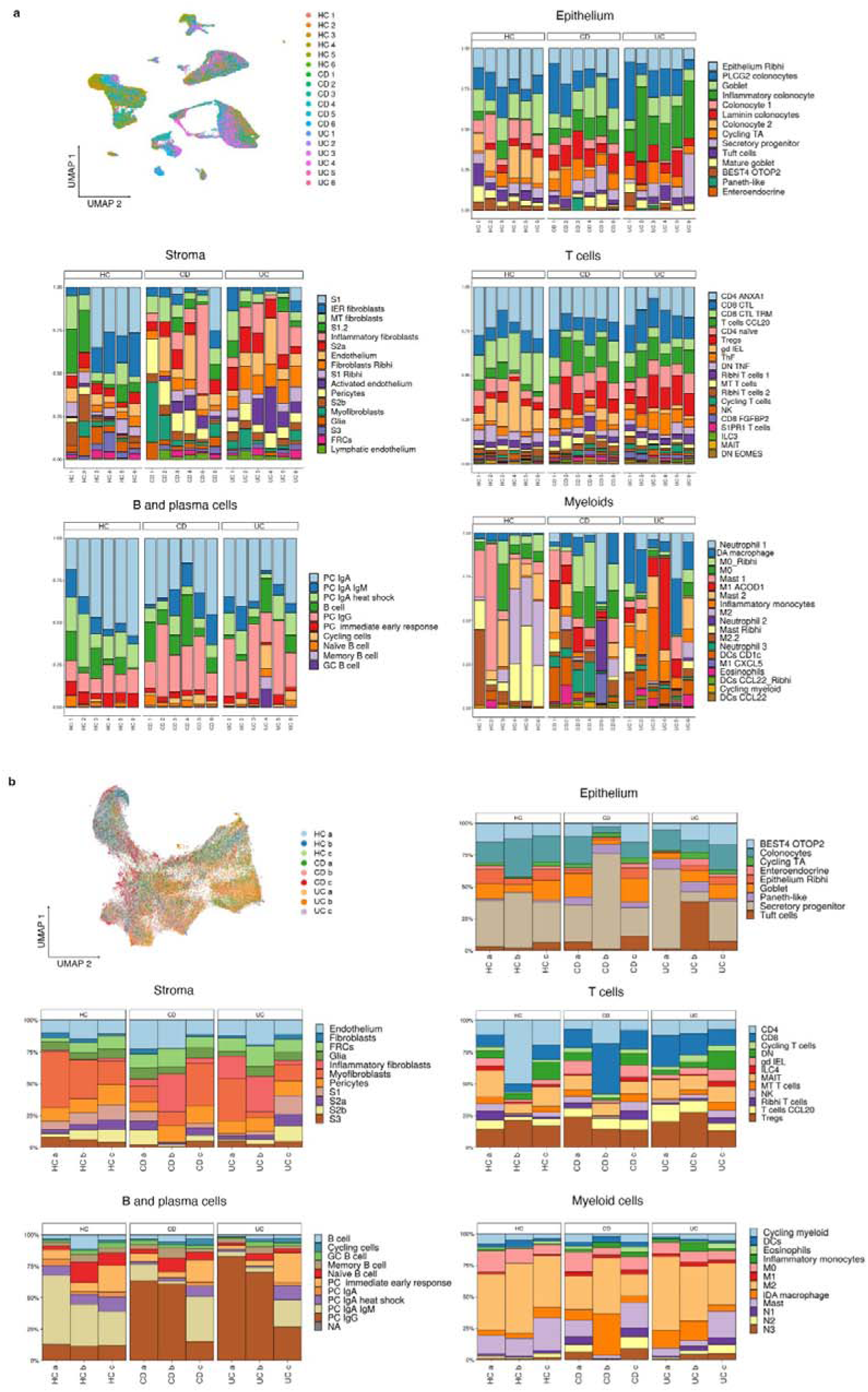
Single-cell RNA-seq (scRNA-seq) and CosMx™ Spatial Molecular Imaging (SMI) cell clusters. **a,** UMAP representation of a sample (n=18) distribution across subsets analyzed by scRNA-seq. Barplots describe the proportions of each cell type within each cell subset (epithelium, stroma, B cells and plasma cells, T cells and myeloid cells) in healthy controls (HC, n=6), active CD (n=6) and active UC (n=6) patients using scRNA-seq data. **b**, UMAP representation of samples analyzed by CosMx™ SMI (n=9). Cells in CosMx™ SMI were annotated by label-transfer of scRNA-seq differentially expressed genes per cell cluster. Barplots describe the proportions of each cell type within each cell subset (epithelium, stroma, B cells and plasma cells, T cells and myeloid cells) in HC (n=3), active CD (n=3) and UC (n=3) patients using CosMx™ SMI data.

**Extended Data Figure 2.**
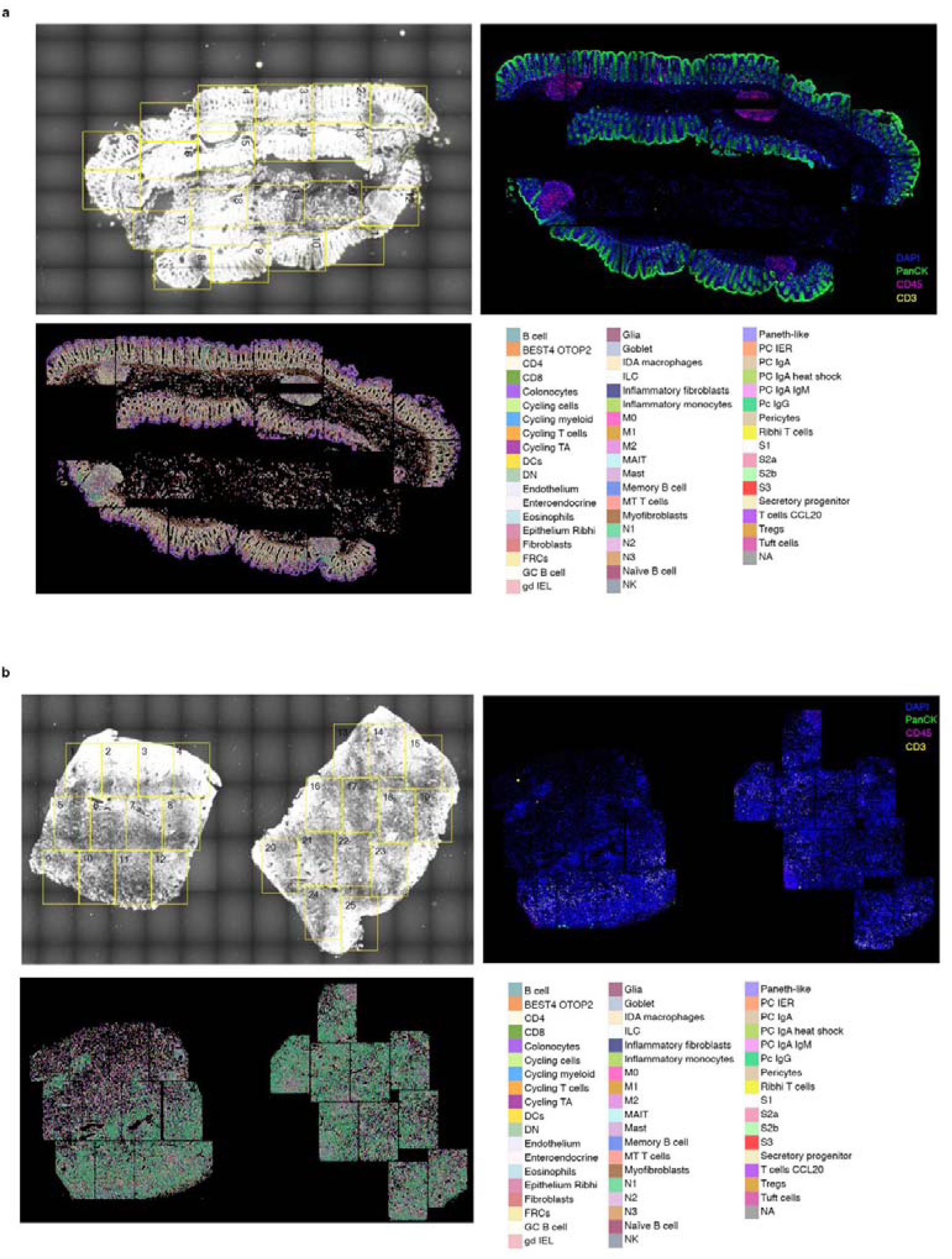
CosMx™ Spatial Molecular Imaging (SMI) of healthy and inflammatory bowel disease (IBD) colonic tissue. Panels show tissue scanner, protein staining and cell type composition of one **a,** healthy control and **b**, ulcerative colitis patient, respectively.

**Extended Figure 3.**
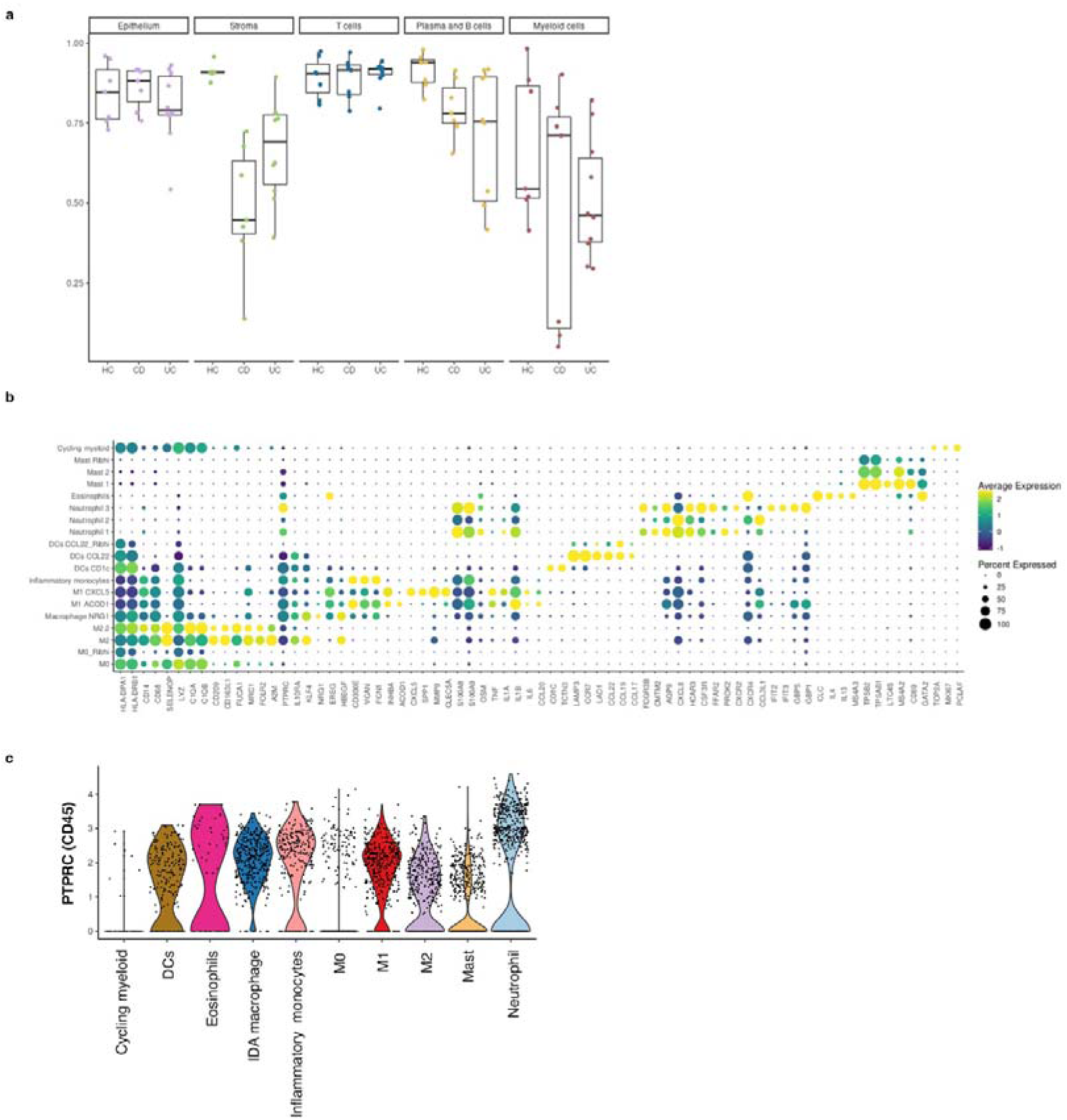
Myeloid characterization. **a**, Morisita-Index analysis of dispersion between samples within each group (HC, CD, UC) for each cell subset (epithelium, stroma, B cells and plasma cells, T cells and myeloid cells) based on scRNA-seq data. **b**, Myeloid cell subsets and their top gene markers. Dot plot shows the fraction of expressing cells (size of the dot) and mean expression levels (dot color). **c**, Violin plot visualization of *PTPRC* (CD45) expression in myeloid populations by scRNA-seq.

**Extended Data Figure 4.**
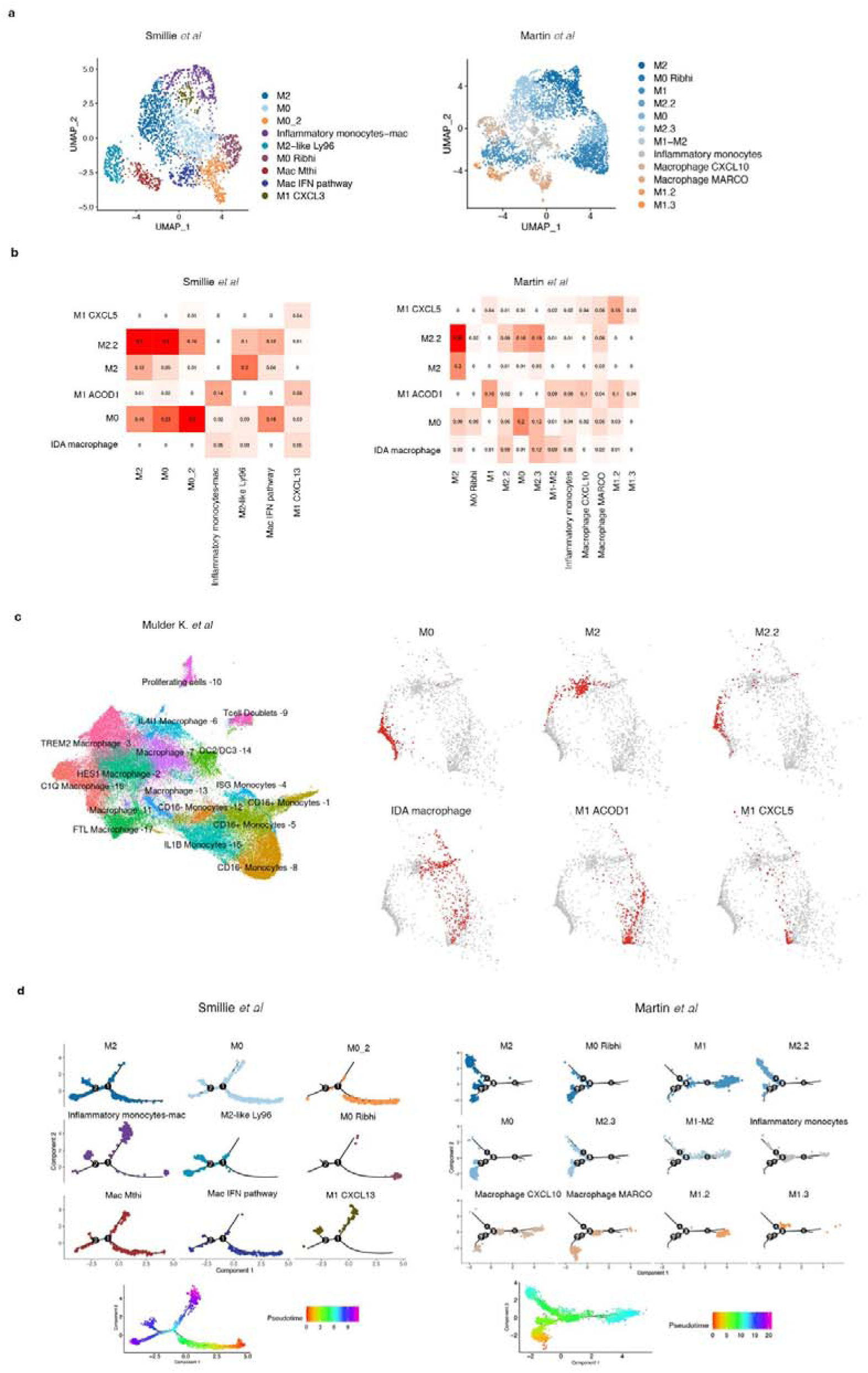
Intestinal Macrophages across studies. **a,** UMAP representation of Smillie *et al-* and Martin *et al-* macrophage subsets isolated *in silico*. **b**, Jaccard similarity index between scRNA-seq data of Smillie *et al-* and Martin *et al-* macrophages and our scRNA-seq macrophage populations. **c**, Projection of our intestinal macrophage clusters found in our study into the MoMAc-VERSE UMAP data (Mulder *et al.2021*). **d**, Pseudo-time trajectory analysis of Smillie *et al-* and Martin *et al- in silico* isolated colonic and ileal macrophages, respectively.

**Extended data Figure 5.**
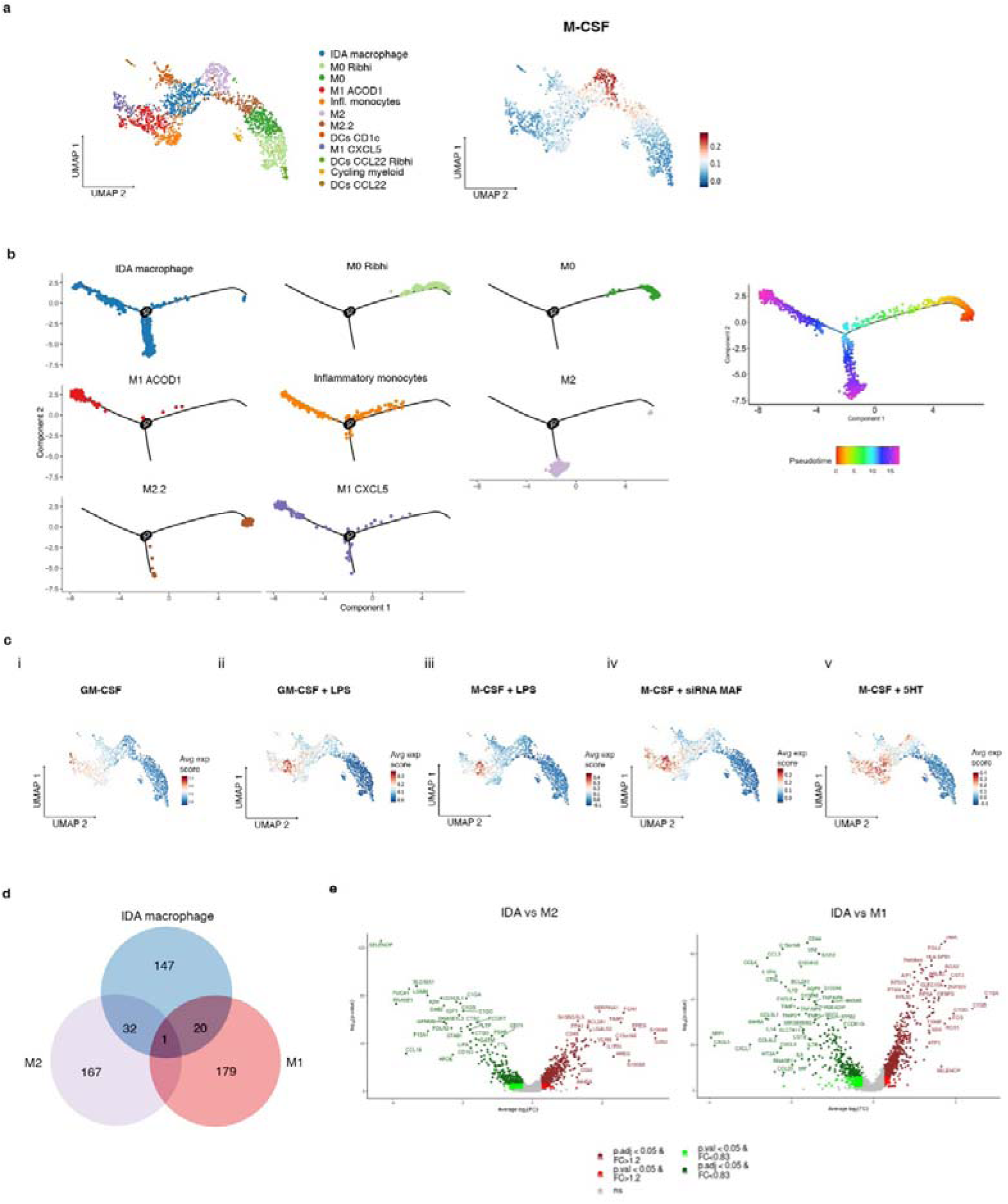
Inflammation-dependent alternative (IDA) macrophages show a unique signature compared to M2 and M1 macrophages. **a,** UMAP showing monocyte, macrophage and dendritic cell clusters identified using scRNA-seq analysis of HC, UC and CD colonic biopsies (left panel). Representation of overlapping signatures (average expression score) of upregulated genes in *in vitro* M-CSF derived macrophages (Cuevas VD *et al*., 2022) on our macrophage cell UMAP (right panel). **b**, Pseudo-time trajectory analysis of monocytes and macrophage populations in our scRNA-seq data. **c**, Representation of overlapping signatures (average expression score) of *in vitro* **i)** GM-CSF-derived macrophages (Cuevas VD *et al*., 2022), **ii)** GM-CSF-derived macrophages stimulated with LPS (Cuevas VD *et al*., 2022), **iii)** M-CSF-derived macrophages stimulated with LPS (Cuevas VD *et al*., 2022), **iv)** upregulated genes of M-CSF-derived macrophages inhibited with MAF siRNA (Vega MA *et al*, 2020) and **v)** upregulated genes of M-CSF-derived macrophages stimulated with 5-HT (serotonin) (Nieto C *et al*, 2020; Domínguez-Soto Á et al, 2017) in our macrophage cell UMAP. **d**, Venn diagram showing the overlap between the top 200 markers of each macrophage population (IDA, M2 (M2 & M2.2) and M1 (M1 ACOD1 & M1 CXCL5) macrophages in the scRNA-seq cohort. **e**, Volcano plot showing differentially expressed genes by scRNA-seq between IDA and M2 or M1 macrophages. Genes upregulated in IDA macrophages are shown in dark (UUP, p value<0.05) or light red (UP, nominal p value<0.05). Genes downregulated in IDA macrophages are shown in dark (DWW, FDR<0.05) or light green (DW, p value<0.05). For each gene, the fold-change (FC) and -log10 (p value) are shown (Supplementary Table 3 contains the complete lists of genes).

**Extended Figure 6.**
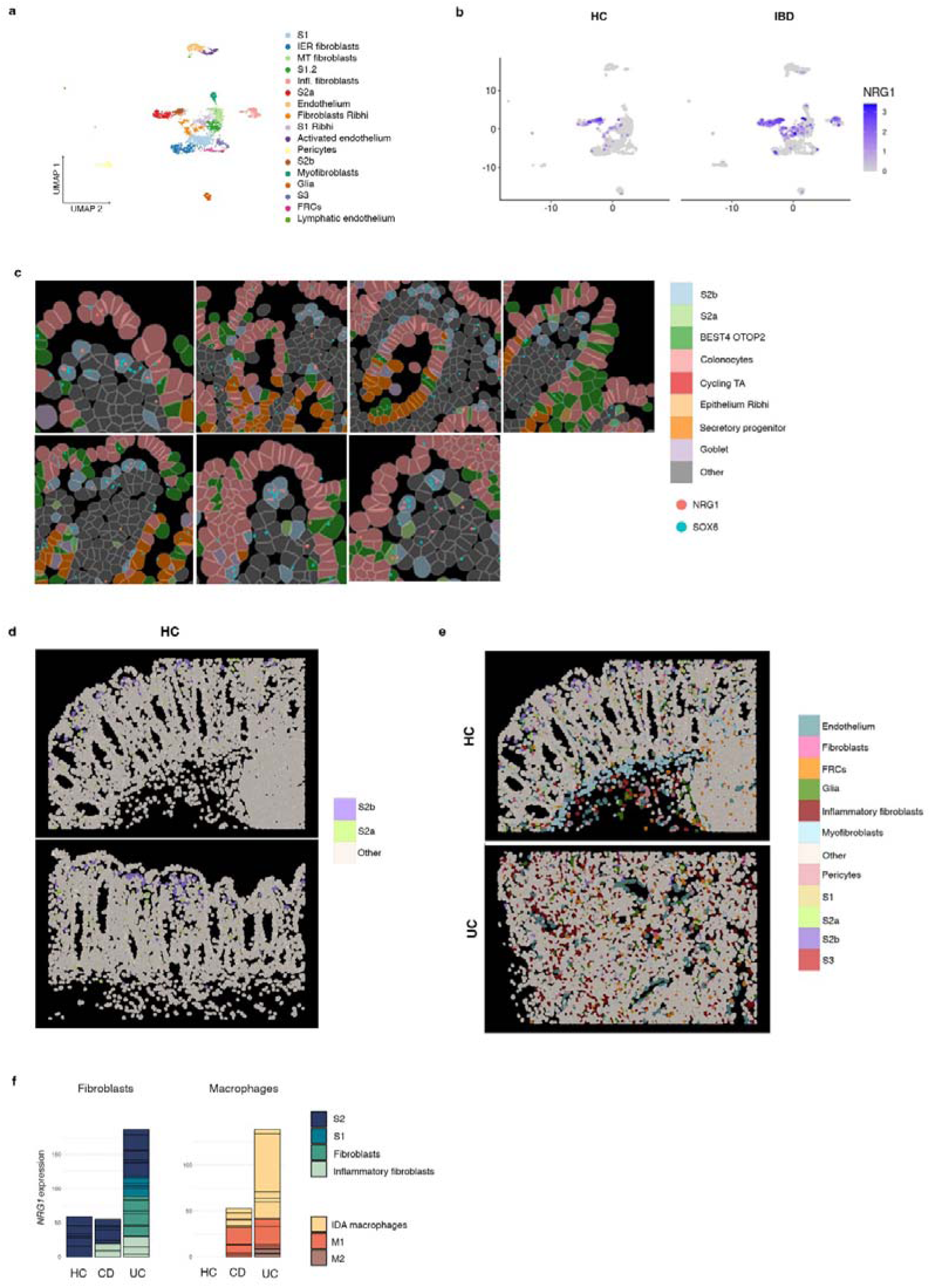
Stromal cell populations analyzed by single-cell RNA-seq (scRNA-seq) and Spatial Molecular Imaging (SMI). **a**, UMAP of stromal clusters observed by scRNA-seq cohort samples. **b**, *NRG1* expression in healthy and IBD stromal subsets in scRNA-seq data. **c**, CosMx™ SMI spatial analysis of S2a and S2b pericryptal fibroblasts showing expression of *NRG1* and *SOX6* (S2 marker) in representative images of a healthy colon. S2b localize at the most apical area. **d**, S2a and S2b differential spatial distribution showed by CosMx™ SMI in 2 representative FoVs from healthy colon. **e**, Spatial localization of stromal cells observed by CosMx™ SMI in representative healthy and inflamed UC colonic mucosa. **f**, *NRG1* expression in the fibroblast and macrophage compartments in HC and IBD colonic tissues according to scRNA-seq data.

**Extended Data Figure 7.**
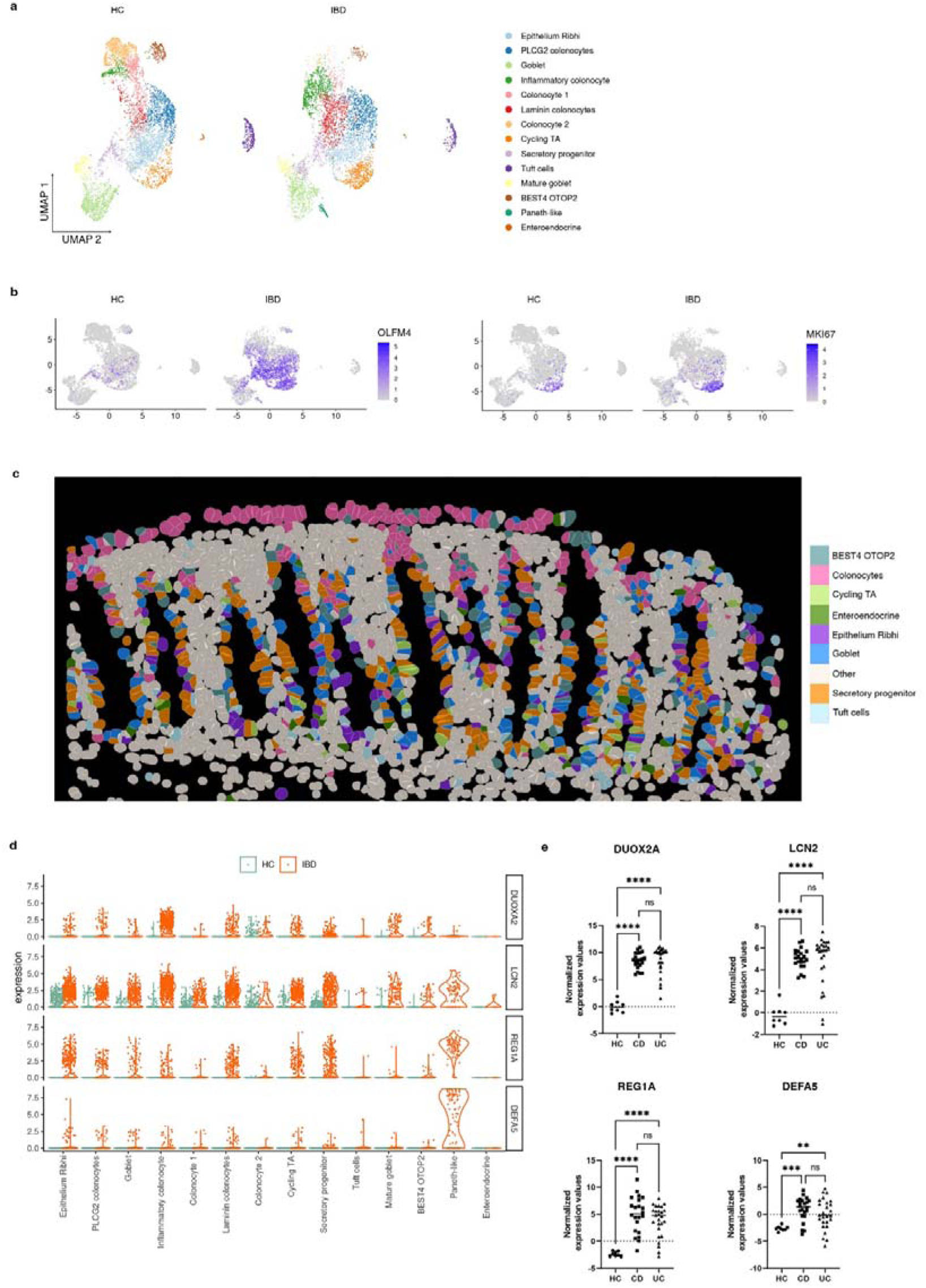
Epithelial cell populations analyzed by single-cell RNA-seq (scRNA-seq) and Spatial Molecular Imaging (SMI). **a**, UMAP of colonic epithelial clusters observed by scRNA-seq in healthy and IBD mucosa (cohort 1). **b**, UMAP showing the expression of *OLFM4* and *MKI67* in the epithelial compartment from healthy control (HC) and IBD tissues (UC and CD) by scRNA-seq. **c**, CosMx™ SMI image of a representative Field of View (FoV) of an HC showing epithelial subsets annotated based on scRNA-seq data. **d**, Violin plots showing the expression (y-axis) of inflammatory markers in epithelial clusters (x-axis) from HC and IBD colons. **e**, Expression of specific inflammatory epithelial cell markers in HC (n=8), active CD (n=22) and active UC (n=26) patients using bulk biopsy RNA-seq data. p<0,05(*), p<0,01 (**), p<0,001(***), p<0,0001(****), ns: not significant.

**Extended Data Figure 8.**
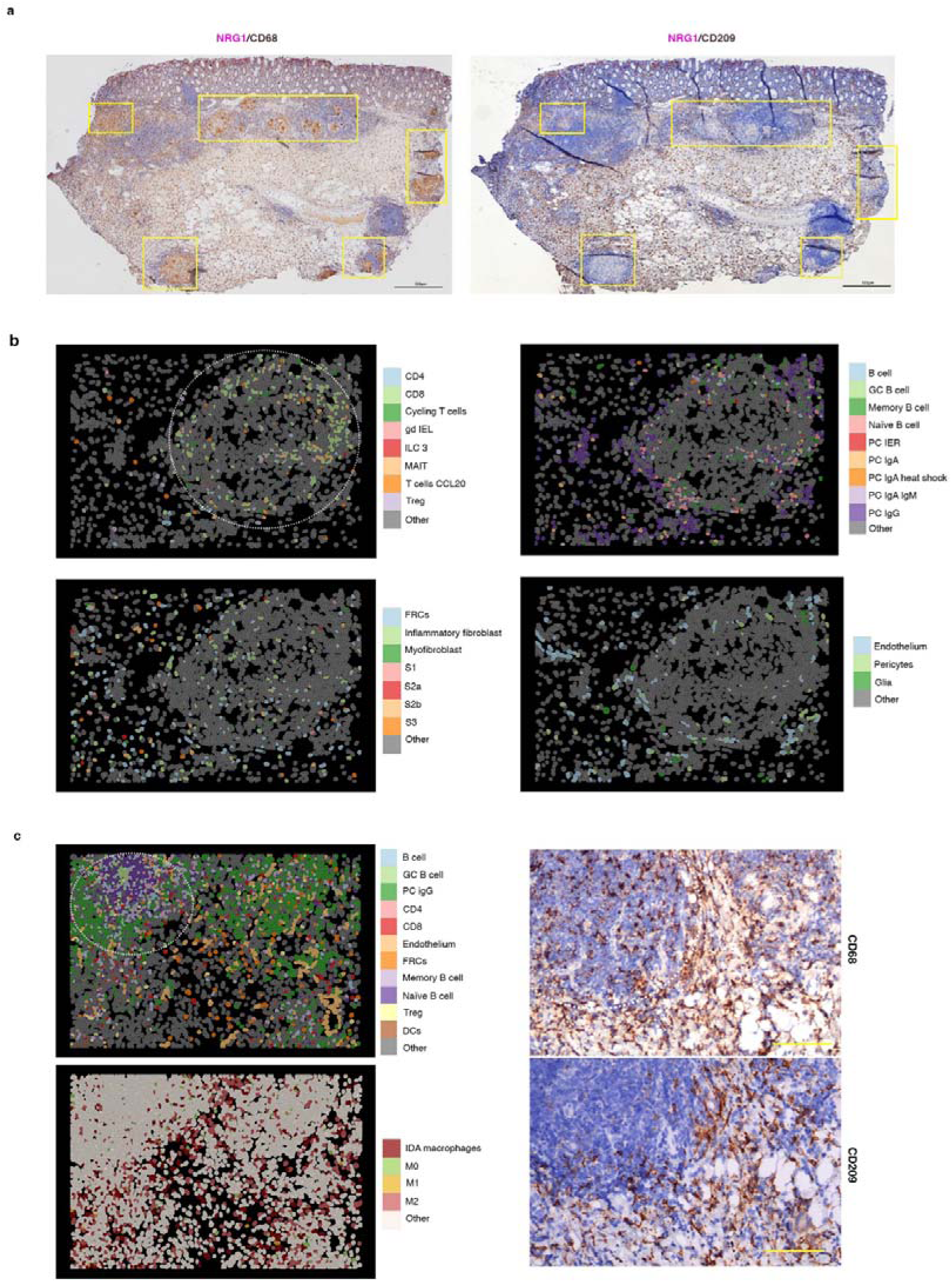
Cellular annotation of all cell types present in Crohn’s disease (CD)- associated granuloma. **a,** Colonic tissue of a CD patient (CD b) containing multiple granulomas indicated by yellow squares. Tissue is stained by *in situ* hybridization for *NRG1* combined with immunostaining for CD68 or CD209. **b**, CosMx™ SMI images showing diverse cell types (stroma cells, T cells, B and plasma cells) present in the granulomas and surrounding area of the analyzed sample. Granuloma is shown by a dotted circle. **c**, Left panels show cell labelling by CosMx™ SMI of a section containing a lymphoid aggregate from the same CD patient. The right panels show expression of CD68 and CD209 by immunohistochemistry in sequential sections (Scale bar= 100 µm). The lymphoid aggregate is shown by dotted circle and is mainly constituted by B cells, abundant plasma cells and some macrophages (M0, M2, M1 and IDA macrophages) are found within the granuloma and more abundantly in the surrounding area.

**Extended Data Figure 9.**
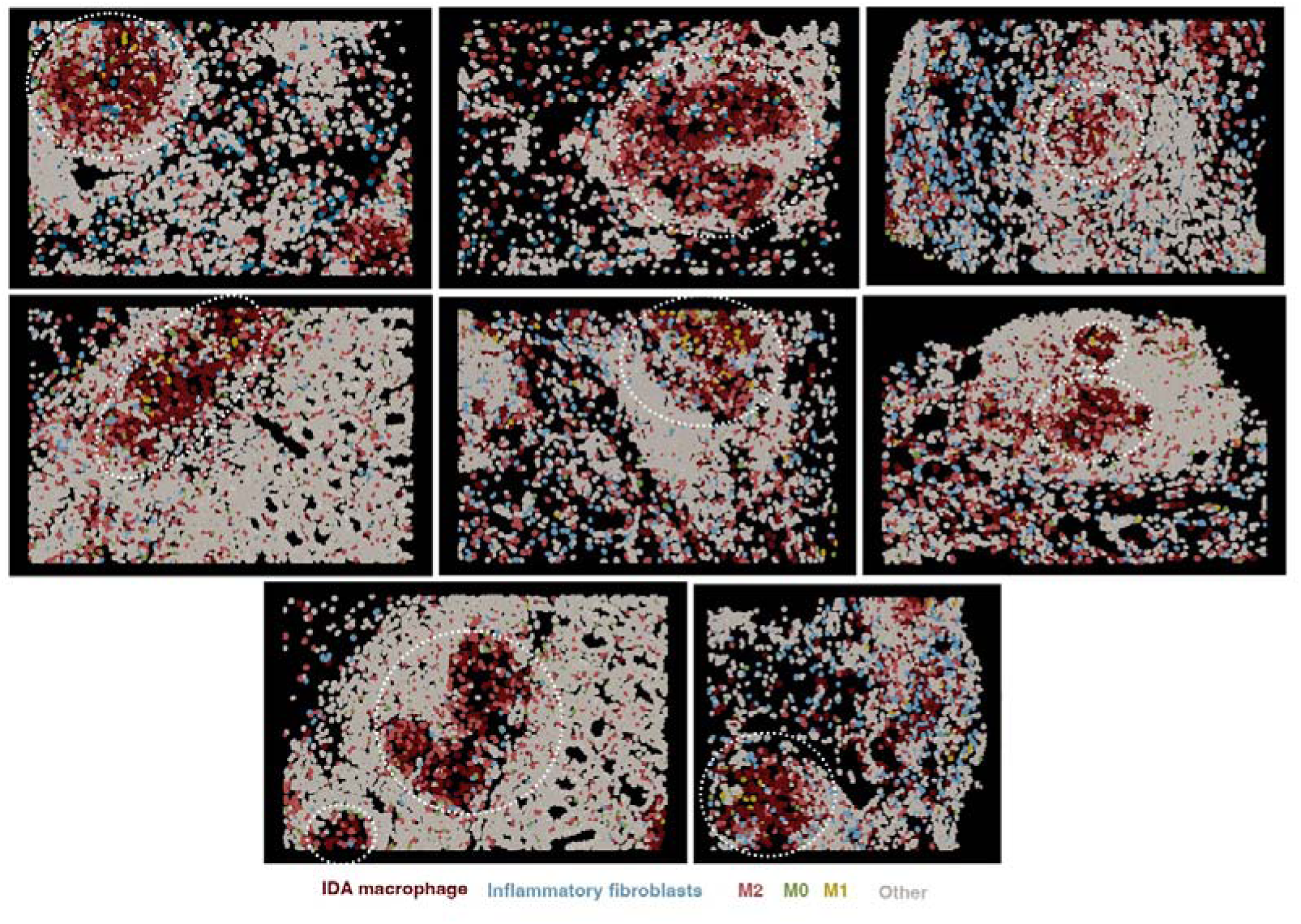
Spatial visualization of inflammatory fibroblasts in inflammatory bowel disease (IBD). **a**, CosMx SMI visualization of inflammatory fibroblasts within granulomas and the surrounding areas in the CD patient that contained multiple granulomas (CD b). Granulomas are indicated by dotted circles. As shown in Figure 4, granulomas contain abundant IDA macrophages. Inflammatory fibroblasts (light blue) can be found within granulomas and in other adjacent areas.

**Extended Data Figure 10.**
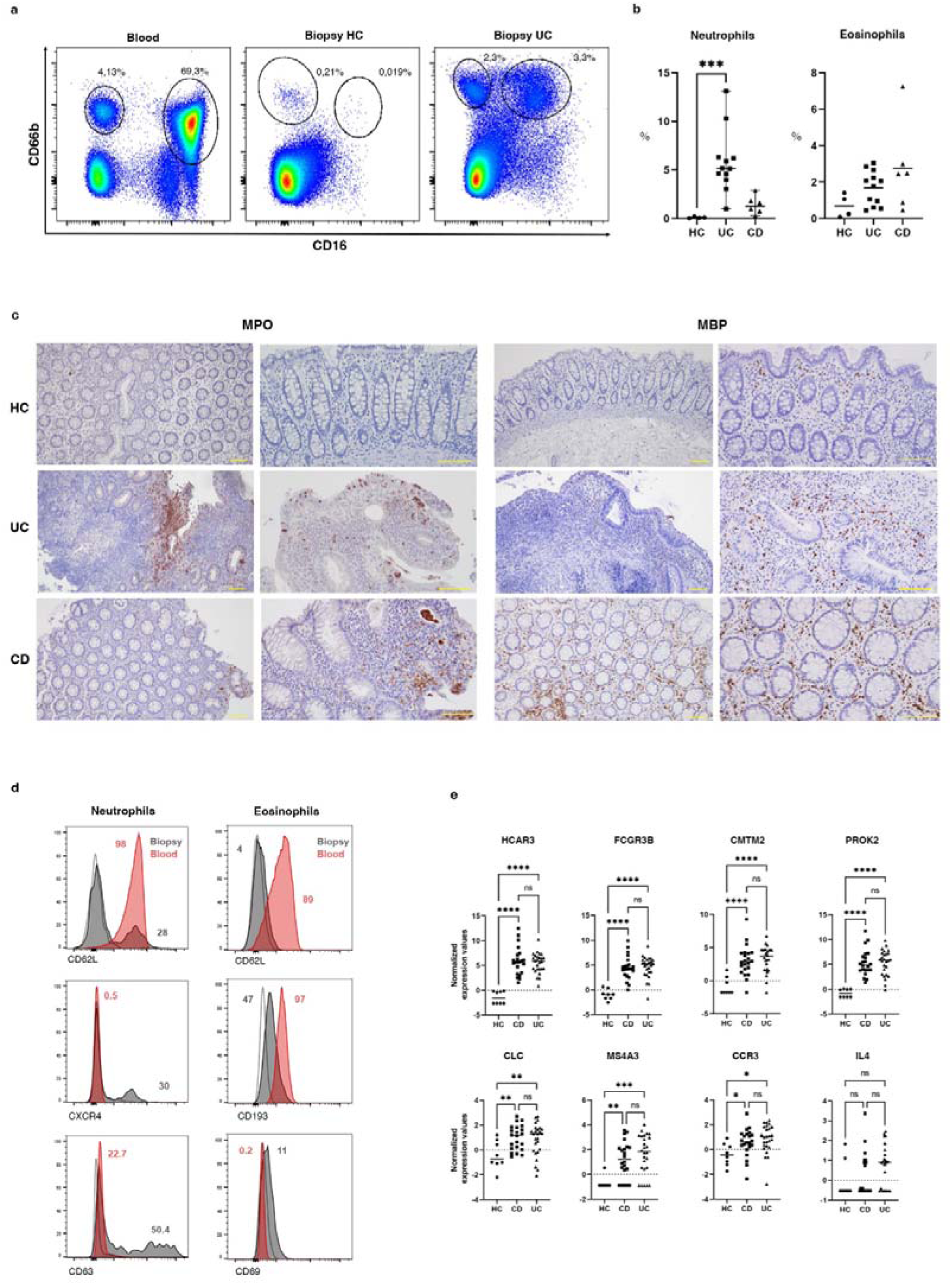
Analysis of neutrophils and eosinophils in inflammatory bowel disease (IBD). **a**, Flow cytometry gating strategy to detect eosinophils and neutrophils in IBD blood and colonic biopsies. Numbers represent the percentages of neutrophils (CD66b^+^ CD16b^+^) and eosinophils (CD66b^+^ CD16b^-^) in the sample displayed as representative of all samples analyzed. **b**, Percentage of neutrophils (CD66b^+^ CD16b^+^) and eosinophils (CD66b^+^ CD16b^-^) in colonic biopsies from healthy controls (HC, n=4), active CD (n=6) and active UC (n=12) colonic samples analyzed by flow cytometry. **c**, Immunostaining for myeloperoxidase (MPO, marker neutrophils) and myelin basic protein (MBP, marker of eosinophils) in representative HC, active UC and active CD colonic tissues (scale bar = 100 µm). **d**, Protein expression detected by flow cytometry in neutrophils (one representative sample is shown from12 colonic and 6 blood samples) and eosinophils (one representative sample shown from 15 colonic and 5 blood samples) from blood and biopsies of IBD patients. Numbers show the percentage of positive cells for the protein in blood (red) and biopsy (grey)Histogram for the corresponding isotype control in the biopsy sample is shown as a dashed line. **e**, Bulk colonic biopsy RNA-seq expression of neutrophil and eosinophil-specific markers in HC (n=8), active CD (n=22) and UC (n=26) patients. p<0,05(*), p<0,01 (**), p<0,001(***), p<0,0001(****), ns: not significant.

