## Supplementary Table 1 for "Macrophage and neutrophil heterogeneity at single-cell spatial resolution in inflammatory bowel disease"

**Supplementary Table 1. Clinical and demographic information of individuals included in the study.**

Endoscopic biopsies (cohort 1) or surgical resection pieces (cohort 2) were processed for single cell RNAseq or spatial molecular imaging (SMI), respectively. Na: non applicable. Nd: no data.

* Score could not be calculated due to incomplete colonoscopy preparation.

|  | Age | Sex | Disease | Biopsy/surgical resection area | Disease location/Extension | Disease duration (years) | Treatment | Total CDEIS | Partial CDEIS | Endoscopic Mayo score |
| --- | --- | --- | --- | --- | --- | --- | --- | --- | --- | --- |
| Cohort 1. scRNA-seq analysis (scRNA-seq) | | | | | | | | | | |
| HC 1 | 61 | Male | na | Sigmoid colon | na | na | na | na | na | na |
| HC 2 | 63 | Male | na | Sigmoid colon | na | na | na | na | na | na |
| HC 3 | 68 | Female | na | Sigmoid colon | na | na | na | na | na | na |
| HC 4 | 65 | Male | na | Sigmoid colon | na | na | na | na | na | na |
| HC 5 | 51 | Male | na | Sigmoid colon | na | na | na | na | na | na |
| HC 6 | 66 | Female | na | Sigmoid colon | na | na | na | na | na | na |
| CD 1 | 49 | Female | CD | Ascending colon | Ileocolonic | 15 | Oral prednisone | 16.61 | 15.5 | na |
| CD 2 | 45 | Male | CD | Sigmoid colon | Ileocolonic | 14 | Adalimumab | 24.24 | 12.1 | na |
| CD 3 | 23 | Female | CD | Descending colon | Ileocolonic | 6 | na | 6.6 | 12 | na |
| CD 4 | 56 | Male | CD | Sigmoid colon | Colonic | 5 | Mesalazine, tacrolimus | 25.4 | 31 | na |
| CD 5 | 21 | Male | CD | Sigmoid colon | Ileocolonic | 5 | Vedolizumab | 13 | 20 | na |
| CD 6 | 22 | Male | CD | Sigmoid colon | Ileocolonic | 10 | Ustekinumab | 4.4 | 9.5 | na |
| UC 1 | 48 | Male | UC | Sigmoid colon | Descending colon | 3 | Oral prednisone, ciprofloxacin, metronidazole | na | na | 3 |
| UC 2 | 43 | Male | UC | Transvers colon | Descending colon | 25 | Mesalazine | na | na | 3 |
| UC 3 | 33 | Female | UC | Rectum | Rectum | 9 | Azathioprine, Prednisone | na | na | 3 |
| UC 4 | 46 | Female | UC | Sigmoid colon | Pancolitis | 5 | Vedolizumab | na | na | 3 |
| UC 5 | 56 | Male | UC | Rectum | Rectum | 6 | Vedolizumab, mesalazine | na | na | 3 |
| UC 6 | 51 | Female | UC | Sigmoid colon | Descending colon | 14 | Mesalazine | na | na | 3 |
| Cohort 2. Spatial molecular Imagining (SMI CosMx, Nanostring) | | | | | | | | | | |
| HC a | 59 | Female | na | Sigmoid colon | na | na | na | na | na | na |
| HC b | 61 | Female | na | Descending colon | na | na | na | na | na | na |
| HC c | 69 | Male | na | Sigmoid colon | na | na | na | na | na | na |
| CD a | 45 | Male | CD | Sigmoid colon | Ileocolonic | 25 | Ustekinumab | * | * | na |
| CD b | 42 | Female | CD | Transverse colon | Colonic | 28 | Cyclosporine | 0 | 8 | na |
| CD c | 38 | Female | CD | Transverse colon | Ileocolonic | 19 | Ciprofloxacin, metronidazole | 14 | 0 | na |
| UC a | 69 | Male | UC | Transverse colon | Pancolitis | 21 | Infliximab | na | na | 3 |
| UC b | 44 | Female | UC | Sigmoid colon | Pancolitis | 5 | Tofacitinib | na | na | 3 |
| UC c | 40 | Male | UC | Sigmoid colon | Pancolitis | 5 | Tofacitinib | na | na | 3 |
| Cohort 3. Bulk RNA-seq | | | | | | | | | | |
| Bulk 1 | 52 | Female | HC | Sigmoid colon | na | na | na | na | na | na |
| Bulk 2 | 51 | Male | HC | Sigmoid colon | na | na | na | na | na | na |
| Bulk 3 | 68 | Male | HC | Sigmoid colon | na | na | na | na | na | na |
| Bulk 4 | 51 | Female | HC | Sigmoid colon | na | na | na | na | na | na |
| Bulk 5 | 27 | Male | HC | Sigmoid colon | na | na | na | na | na | na |
| Bulk 6 | 49 | Female | HC | Sigmoid colon | na | na | na | na | na | na |
| Bulk 7 | 32 | Female | HC | Sigmoid colon | na | na | na | na | na | na |
| Bulk 8 | 52 | Female | HC | Sigmoid colon | na | na | na | na | na | na |
| Bulk 9 | 33 | Male | CD | Sigmoid colon | Colonic | 13 | Immunosupressor | 10.4 | 16 | na |
| Bulk 10 | 47 | Male | CD | Descending colon | Colonic | 24 | Immunosupressor | 19 | 20 | na |
| Bulk 11 | 53 | Female | CD | Rectum | Ileocolonic | 28 | No treatment | 20.5 | 34 | na |
| Bulk 12 | 39 | Female | CD | Descending colon | Ileocolonic | 23 | Immunosupressor | 6.2 | 31 | na |
| Bulk 13 | 67 | Male | CD | Sigmoid colon | Colonic | 7 | Steroids | 12 | 30 | na |
| Bulk 14 | 35 | Female | CD | Descending colon | Ileocolonic | 7 | No treatment | 17.75 | 20.5 | na |
| Bulk 15 | 64 | Female | CD | Sigmoid colon | Ileocolonic | 7 | Steroids | 19.75 | 31 | na |
| Bulk 16 | 35 | Male | CD | Ascending colon | Ileocolonic | 27 | Immunosupressor | 12.6 | 12 | na |
| Bulk 17 | 31 | Female | CD | Descending colon | Colonic | 6 | No treatment | 13.5 | 19 | na |
| Bulk 18 | 33 | Male | CD | Sigmoid colon | Colonic | 21 | Immunosupressor | 6.8 | 17 | na |
| Bulk 19 | 56 | Female | CD | Sigmoid colon | Colonic | 14 | ND | 7.88 | 9.5 | na |
| Bulk 20 | 22 | Male | CD | Descending colon | Colonic | 9 | Immunosupressor | 22.6 | 32 | na |
| Bulk 21 | 42 | Male | CD | Transverse colon | Ileocolonic | 25 | ND | 4.67 | 5 | na |
| Bulk 22 | 56 | Female | CD | Rectum | Ileocolonic | 29 | ND | 15.88 | 29.5 | na |
| Bulk 23 | 44 | Female | CD | Rectum | Colonic | 12 | Immunosupressor | 10.4 | 26 | na |
| Bulk 24 | 37 | Male | CD | Ascending colon | Ileocolonic | 23 | Immunosupressor | 24 | 30 | na |
| Bulk 25 | 20 | Male | CD | Rectum | Colonic | 14 | Immunosupressor | 12.8 | 32 | na |
| Bulk 26 | 24 | Female | CD | Sigmoid colon | Colonic | 9 | Steroids | 5.6 | 10 | na |
| Bulk 27 | 60 | Female | CD | Transverse colon | Colonic | 33 | Immunosupressor | 4.67 | 14 | na |
| Bulk 28 | 29 | Male | CD | Ascending colon | Colonic | 11 | Immunosupressor | 19.6 | 17 | na |
| Bulk 29 | 48 | Male | CD | Sigmoid colon | Ileocolonic | 32 | Immunosupressor | 35.5 | 34 | na |
| Bulk 30 | 41 | Male | CD | Rectum | Colonic | 16 | Immunosupressor | 8.25 | 16.5 | na |
| Bulk 31 | 57 | Female | UC | Sigmoid colon | Pancolitis | 11 | Immunosupressor | na | na | 3 |
| Bulk 32 | 43 | Male | UC | Sigmoid colon | Pancolitis | 30 | Immunosupressor, Steroids | na | na | 2 |
| Bulk 33 | 56 | Male | UC | Sigmoid colon | Pancolitis | 25 | Immunosupressor | na | na | 3 |
| Bulk 34 | 61 | Female | UC | Descending colon | Left-sided colitis | 38 | Immunosupressor | na | na | 3 |
| Bulk 35 | 31 | Male | UC | Sigmoid colon | Pancolitis | 9 | Immunosupressor, Mesalazine | na | na | 3 |
| Bulk 36 | 62 | Male | UC | Sigmoid colon | Left-sided colitis | 13 | Steroids | na | na | 2 |
| Bulk 37 | 44 | Male | UC | Sigmoid colon | Pancolitis | 24 | Immunosupressor | na | na | 3 |
| Bulk 38 | 28 | Male | UC | Sigmoid colon | Pancolitis | 7 | Immunosupressor | na | na | 3 |
| Bulk 39 | 29 | Female | UC | Sigmoid colon | Pancolitis | 10 | Immunosupressor, Steroids | na | na | 3 |
| Bulk 40 | 55 | Male | UC | Sigmoid colon | Pancolitis | 30 | Immunosupressor | na | na | 3 |
| Bulk 41 | 48 | Male | UC | Rectum | Proctitis | 37 | Steroids | na | na | 2 |
| Bulk 42 | 37 | Female | UC | Rectum | proctitis | 25 | ND | na | na | 2 |
| Bulk 43 | 19 | Male | UC | Ascending colon | Pancolitis | 9 | ND | na | na | 2 |
| Bulk 44 | 31 | Male | UC | Sigmoid colon | Pancolitis | 11 | Immunosupressor | na | na | 3 |
| Bulk 45 | 37 | Female | UC | Rectum | Left-sided colitis | 14 | Immunosupressor, Steroids | na | na | 3 |
| Bulk 46 | 35 | Male | UC | Sigmoid colon | Left-sided colitis | 14 | Steroids, Mesalazine | na | na | 3 |
| Bulk 47 | 69 | Male | UC | Sigmoid colon | Left-sided colitis | 12 | Immunosupressor | na | na | 3 |
| Bulk 48 | 25 | Female | UC | Sigmoid colon | Pancolitis | 10 | Immunosupressor | na | na | 2 |
| Bulk 49 | 38 | Male | UC | Sigmoid colon | Left-sided colitis | 14 | Immunosupressor | na | na | 3 |
| Bulk 50 | 50 | Male | UC | Sigmoid colon | Pancolitis | 29 | Mesalazine | na | na | 3 |
| Bulk 51 | 41 | Male | UC | Sigmoid colon | proctitis | 14 | No treatment | na | na | 3 |
| Bulk 52 | 39 | Female | UC | Sigmoid colon | Left-sided colitis | 22 | No treatment | na | na | 3 |
| Bulk 53 | 50 | Male | UC | Sigmoid colon | na | 29 | No treatment | na | na | na |
| Bulk 54 | 40 | Female | UC | Sigmoid colon | na | 22 | Stop anti-TNF (Golimumab) | na | na | na |
| Bulk 55 | 59 | Female | UC | Rectum | Ileocolonic | 31 | Immunosupressor | na | na | na |
| Bulk 56 | 27 | Female | UC | Sigmoid colon | Colonic | 7 | Immunosupressor | na | na | na |
