## Supplementary material for "Macrophage and neutrophil heterogeneity at single-cell spatial resolution in inflammatory bowel disease": Methods

**Patient recruitment and sample collection**

Colon samples were collected to perform scRNAseq, generate organoid cultures and analyze by flow cytometry on fresh tissue. Additional samples were fixed in formalin and paraffin embedded (FFPE) for CosMx^TM^ SMI and tissue staining (IHC IF, ISH). For scRNAseq and flow cytometry, colon biopsies from active areas of UC and CD patients were collected during routine endoscopies performed as standard of care. Healthy controls were individuals undergoing endoscopy for colorectal cancer screening and presenting no signs of dysplasia or polyps at the time of endoscopy. For SMI analysis, surgical colon resections were obtained from non-IBD controls (patients undergoing surgery for colorectal cancer), UC and CD patients (undergoing colonic resective surgery). Organoid cultures were established exclusively from surgical samples of non-IBD controls. In non-IBD controls samples were obtained from non-tumor areas and for UC and CD patients from involved inflamed areas. Blood samples from HC, CD and UC were also collected to perform flow cytometry analysis of granulocytes. The study was approved by the Ethics Committee of Hospital Clinic Barcelona (HCB/2018/1062 and HCB/2022/0125) and the Hospital Mutua de Terrassa (CI201901). All patients signed an informed consent at the time of colonoscopy or before surgical intervention.

**Human colonic cell isolation**

Biopsies (n=4-6 per patient) were taken from involved areas of the colon of UC and CD patients with signs of endoscopic activity, placed immediately in cold Hank’s Balanced Salt Solution (HBSS) (Gibco, MA, USA) and kept at 4ºC until processing (<1 h). Colonic biopsies from non-IBD controls were collected from the sigmoid colon and processed in the same way. Freshly collected biopsies were washed with 5mM DTT (Roche, Spain) in HBSS for 15 min and then washed in complete medium (CM) (RPMI 1640 medium (Lonza, MD, USA) supplemented with 10% heat-inactivated fetal bovine serum (FBS) (Biosera, France), 100U/ml penicillin, 100 U/ml streptomycin and 250 ng/ml amphotericin B (Lonza), 10µg/ml gentamicin sulfate (Lonza) and 1,5mM Hepes (Lonza)) for 10 minutes. Both incubations were performed at room temperature in a platform rocker. Biopsies were chopped with a scalpel and placed into tubes containing 500 μl of Digestion Solution (CM + Liberase TM (0.5 Wünsch units/ml) (Roche, Spain) + DNase I (10 μg/mL) (Roche, Spain)) and incubated on a shaking platform for 1h at 250 RPM and 37ºC. After incubation biopsies were filtered through a 50-μm cell strainer (CellTrics, Sysmex, USA), washed with Dulbecco’s Phosphate Buffered Saline (PBS; Gibco, USA) and resuspended in RPMI medium supplemented with 0.05% of Bovine serum albumin (BSA) at a concentration of ~0.5-1·10^6^ cells/mL for scRNA-seq and in FACS buffer (PBS + 2% inactivated FBS (fetal bovine serum) + NaN3 0.1%) for flow cytometry analysis ^1^

**10x library preparation and sequencing**

Following digestion, 10x Genomics 3′ mRNA single-cell method was used. Approximately 7,000 cells were loaded onto the Chromium10x Genomics platform (10x Genomics, CA, USA) to capture single cells, following the manufacturer’s protocol. Generation of gel beads in emulsion (GEMs) (10x Genomics, CA, EEU), barcoding and GEM-reverse transcription was performed using the Chromium Single Cell 3′ and Chromium Single Cell V(D)J Reagent Kits (10x Genomics, CA, EEU) (user guide, no. CG000086) according to manufacturer’s instructions. Full-length, barcoded cDNA was amplified by PCR to generate enough mass for library construction (Nextera® PCR primers) (Illumina, CA, USA). Sequencing of the libraries was performed on HiSeq2500 (Illumina, CA, USA).

**Single cell data analysis**

***Data Processing***

For each sample, sequences obtained in fastq files were processed with CellRanger’s count pipeline using the default parameters (10XGenomics, v. 3.1.0). This pipeline performs an alignment based on the reference genome (Gencode release 27, assembly GRCh38 p10), filtering, barcode counting, and UMI counting. The resulting filtered matrix was analyzed using R (v.4.2.0). We merged the count matrices retrieved from CellRanger using the function *merge* from the *SeuratObject* R package (version 4.0.2). At this point, doublets were assessed using the *scDblFinder ^2^*R package and removed. Total analyzed cells: 47,600.

Initially, we analyzed healthy control and IBD samples separately to assess the similarity of cell types and samples. We processed and annotated the objects separately and we assessed for similarity by using Jaccard index and label transferring (see detailed methods in [here](https://servidor2-ciberehd.upc.es/external/garrido/methods1/)). Samples were then pooled together in the same object. Low-quality cells were then filtered out based on mitochondrial RNA percentage and number of genes per cell. Epithelial cells required a less stringent filter of 65% of counts aligned to the mitochondrial genes for quality control. A total of 46,700 cells were considered for the analysis. Then, we logarithmically normalized, obtained the highly variable genes, and scaled the counts (default parameters) of each data set using *Seurat ^3^*(version 4.1.0). Principal component analysis (PCA) was performed. Dimensionality reduction was performed by applying the Uniform Manifold Approximation and Projection (UMAP) algorithm using the optimal number of PCs ^4^. UMAP also served as a two-dimensional embedding for data visualization. Cluster analysis was performed using the Louvain clustering algorithm^3^

We used marker gene expression to categorize and split cells in the main five subsets to analyze them separately. *EPCAM, AQP8, BEST4, MUC2, OLFM4, PLCG2, TRPM5, ZG16 for epithelial cells; CD79A, BANK1, CD19, DERL3, MS4A1, MZB1* for B and cells; C*D3D, CD3E, CD3G, CD8A, FOXP3, GZMA, GZMB, IL17A, NKG7, TRBC1* for T cells; *ACTA2, ADAMDEC1, CHI3L1, COL3A1, NRXN1, PVALP, SOX6, VWF* for stromal cells; *AIF1, C1QA, C1QB, CD14, CMTM2, FCGR3B, LYZ, MS4A2, TPSAB1, TPSAB2* for myeloid cells. We then separated the merged data into 5 main cell types: Epithelial cells, Lymphoid cells (T cells and innate lymphoid cells), Myeloid cells (mast cells, macrophages, dendritic cells, monocytes, neutrophils, and eosinophils), Stromal cells (endothelial cells, fibroblasts, pericytes and glia), and B and Plasma cells. Immunoglobulin (IG) genes were removed from all the main cell types except B and plasma cells to reduce background noise.

***Identification of cell types***

Each main cell type was re-processed starting from *FindVariableFeatures* using the same procedure as for the whole dataset (see full code in our [github page](https://gitfront.io/r/user-1871750/StAFev2VqZh6/ibd-bcn-single-cell/) and our [webpage](https://servidor2-ciberehd.upc.es/garrido.html)). During the process, we further removed *CD3*, *C1QA*, *DERL3*, or *MS4A1* expressing cells from Epithelial cells; *CD3, C1QA*, or EPCAM-expressing cells from the B cells; *CD3*, *C1QA*, *DERL3*, *MS4A1* or *EPCAM*-expressing cells from the Stromal cells; *DERL3, MS4A1, C1QA, EPCAM,* or *CD79A*-expressing cells from the Lymphoid cells; and *CD3, THY1, DERL3* or *MS4A1*-expressing cells from the Myeloid cells. We then systematically re-clustered each cell category. The annotation of each subcluster from the main cell type was defined by the marker genes, which were obtained by the *FindAllMarkers* function with the default threshold except for the *min.pct* parameter, which was set as 0.25, and the *thresh.use*, which was set to 0.25.

***Batch correction***

We observed batch effect by sample in the UMAPs of Lymphoid (tcells), B and plasmacells (plasmas), and Stromal cells (stroma) objects while re-processing for cell type identification. For those cell types, we ran *Harmony ^5^*(version 0.1.0) to correct for this effect. Specifically, we used the RunHarmony function, with the optimal number of principal components found for each subset as latent space and the sample of origin as batch label. *Seurat’s* analysis pipeline was applied again to each object, followed by UMAP generation using default settings and harmony integrated space.

***Annotation of cells***

*Epithelial cell subset (EPCAM positive)*: Within the absorptive cells, we found two clusters of colonocytes sharing the expression of *AQP8, FABP1* and *SLC26A2*, and a cluster of inflammatory colonocytes with low *AQP8* expression and expressing the inflammatory markers *DUOXA2, DUOX2* and *LCN2*. PLCG2 colonocytes express *PLCG2, HES1* and *ELF1*, while the Laminin enterocytes present expression of the laminin gene *LAMA3* and the Ephrin receptor *EPHA2*. Last, a cluster of BEST4 colonocytes (*BEST4* and *OTOP2*) and Tuft cells (*TRPM5*, *SH2D6* and *POU2F3*) were identified. Within the secretory cells, we found two clusters of goblet cells (*MUC2, TFF3* and *SPINK4*), a canonical and a mature cluster expressing *TFF1* and *FER1L6*. Secretory progenitor cells express *RETNLB, CLCA4* and *HEPACAM2*. A cluster of Paneth-like cells was identified in the inflamed samples expressing the defensin genes *DEFA5* and *DEFA6* and *REG3A*. Enteroendocrine cells were identified by *CHGA*, *CHGB* and *NEUROD1*. Cells with high expression of ribosomal genes (*RPS19, RPS18, RPL35*) clustered together and were annotated as Epithelium Ribhi. While *LGR5*+ stem cells did not form a cluster, we identified cycling transit-amplifying cells by the expression of *MKI67, TOP2A* and *PCNA*.

*Stromal cell subset*: fibroblasts were annotated based on the nomenclature of Kinchen et al^6^. Three clusters of S1 lamina propria fibroblasts were identified, all sharing the expression of *ADAMDEC1*, *CP* and *FABP4*. Submucosal S3 fibroblasts express *OGN*, *CCDC80* and *GREM1*. Peri-cryptal S2 (*SOX6*, *F3*) fibroblasts subclustered into the bottom crypt S2a (*VSTM2*) and the top-crypt S2b (*NPY, NRG1*) clusters. Moreover, we also found a cluster of Rib^hi^ fibroblasts (*RPL7A, RPL28*), a cluster of MT fibroblasts (with high expression of mitochondrial genes) and an IER fibroblasts cluster, characterized by the expression of immediate-early response genes (*FOS, FOSB, IRF1*). Fibroblastic reticular cells, FRCs, were identified by the expression of *CCL19* and *CCL21,* and myofibroblasts by the expression of *SOSTDC1, ACTG2* and *MYH11*. Last, inflammatory fibroblasts (also known as S4) were identified by the expression of *IL11, CXCL5, FAP, INHBA* and *IL24*, among others.

Endothelial cells express *VWF, PECAM1* and *PLVAP*. In inflammation, a cluster of activated endothelium was identified expressing *ACKR1* and the selectins *SELE* and *SELP*. We further identified a small lymphatic endothelium cluster expressing *MRC1*, *MMRN1* and *LYVE1*.

Finally, pericytes were characterized by the expression of *NOTCH3* and *RGS5;* and the markers *NRXN1*, *S100B* and *CDH19* identified a cluster of glial cells.

*B and plasma cell subset*: Plasma cells (*DERL3, MZB1, XBP1*) could be subdivided based on whether they are IgA, IgG or IgM producers, and whether they are plasmablasts, PB, (PRDM1^low^) or fully differentiated plasma cells, PC (PRDM1^high^)^7,8^. We found some clusters expressing in an exclusive manner *IGLC3* and *IGLC2*, which we classified as a Lambda cluster, and a plasma cell cluster expressing *IGLL5, IGLC7* and *IGLL1*, annotated as PC IGLL5. Moreover, two clusters of IgA plasma cells expressing the heat-shock genes *HSPA1B, HSPA1* and *HSPB1* were annotated as PC IgA heat shock. As in other subsets, cells expressing immediate-early genes (*FOS*, *JUN* and *FOSB*) were annotated as PC IER.

Unlike plasma cells which loss its expression through differentiation, B cells were identified by the expression of CD20 (*MS4A1*). Within the B cell lineage, we identified a cluster of memory B cells (*ITGAX*, *FGR* and *PDCD1*), naive B cells (*IGHD* and *FCER2*) and germinal center (GC) B cells (*SUGCT, MME* and *HRK*). Besides, B cells with high expression of ribosomal genes were annotated as B cell Ribhi. Last, two clusters of cells in a proliferative state were identified, annotated as cycling cells (*TUBB, TOP2A*, *MKI67*) and including a mixture of B and plasma cells.

*T cell subset*: T cells were determined by the expression of T cell receptor component *CD3D, CD3E* and *CD3G*. CD8 T cells had expression of *CD8A, CD8B,* or both, and CD4 cells were *CD4^low^.* Withing the CD4 cells, CD4 naïve cells were identified by the expression of *CCR7*, *LEF1* and *SELL*. CD4 ANXA1^9^ cluster express *ANXA1*, *IL7R* and *GPR186,* and could represent a resident phenotype. Tregs were identified by the expression of *FOXP3*, *IL2RA*, *CXCR6* and *BATF,* among others. T follicular helper cells, ThF, express *CXCL13* and *MAGEH1*. While we did not observe a canonical Th17 population, the CCL20 cluster represent a heterogeneous population with inflammatory related genes that probably include Th17 cells (*CCL20*, *RORA*, *KLRB1*, *ETPD1*)^10^. S1PR1 T cells correspond to a circulating phenotype (Central Memory Cells) and express *CCR4*, *KLF2* and *S1PR1* among others. Within the CD8 cells, CD8 cytotoxic effector cells were identified by the expression of *GZMK* and *KLRG1,* while CD8 TRM lack expression of GZMK but express other GZM genes, KLRB*1, SPRY1* and *IGTA1*. CD8 FGFBP2 corresponds to cytotoxic cells described previously in ulcerative colitis^11^, which expresses *FGFBP2*, *FCGR3A*, *S1PR5* along with cytotoxic genes (*GZMH*, *NKG7*). The MAIT cluster is *CD8A*^+^ and were identified by the expression of *NRC3*. DN TNF is a cluster of double negative T cells (*CD8*^-^*CD4*^-^) expressing immediate-early response genes (*FOS*, *JUN* and *FOSB*) phenotype and the inflammatory cytokine *TNF*. DN EOMES is a cytotoxic effector DN cluster (*CD8*-*CD4*-) characterized by the expression of the transcription factor *EOMES* and *GZMK.* GD IEL correspond to a cluster of intraepithelial lymphocytes with expression of the T receptor genes *TRDC* and *TRGC1*, which is further identified by the expression of *FCER1G*, *KLRC2*, *CD160*, *KLRD1*, *KLRC1*, *ITGA1* and *KIR2DL4^12^*. Regarding the innate lymphocytes, we observed a cluster of ILC3 (*LIF*, *KIT* and *PCDH9)* and NK (expressing *KLRF1* and lacking expression of *CD3G*, *CD3D*, *CD8A* and *CD4)*.

*Myeloid cell subset*: The myeloid cell subset includes macrophages, monocytes, dendritic cells, mast cells eosinophils and neutrophils. Macrophages were identified by the expression of *CD68, CD14* and complement coding-genes *(C1QA, C1QB)*. While M2 and M2.2 macrophages express *CD209*, *CD163L1* and *FOLR2,* M0 macrophages lack the expression of M2 markers but express macrophage marker genes (*CD68*, *C1QA*, *C1QB* and so on). The IDA subset was identified by the expression EGF ligands (*NRG1 AREG*, *EREG* and *HBEGF*). M1 ACOD1 and M1 CXCL5 shared the expression of *TNIP3*, *IL1B*, *INHBA*, *IL6*, *VCAN*, *CD300E* but express *ACOD1* and *CXCL5*, respectively, in an exclusive manner^13^. Unlike M1, inflammatory monocytes were identified by higher expression of *VCAN, CD300E* and *CD14*, and the expression of *FCN1 and* *S100A9.* We identified two clusters of mast cells (*TPSB2* and *TPSAB1),* one of them expressing *LTC4S.* We further identified eosinophils by the expression of CLC, *IL4* and *IL13*. All neutrophil clusters were identified by the expression of *PROK2*, *CMTM2*, *CXCL8, FCRG3B, AQP9,* *S100A8* and *S100A9*. Finally, two clusters of dendritic cells were identified (*LAMP3)*. On one side, DCs CD1c express *CD1c*, *TRL10* and *TCTN3,* while DCs CCL22 were identified by the expression of *CCL22* and *CCL19*.

#### ***Differential abundance testing***

We tested for differences in cell abundances between myeloid colonic cells from IBD patients and healthy controls using the *miloR* package (v 1.4.0)^14^. Specifically, we performed differential abundance testing of healthy patients and both inflammatory chronic diseases, CD and UC, separately. To do so, the dataset was subsetted to include myeloid cells only (677 healthy cells, 1935 CD cells and 1161 UC myeloid cells). For each analysis, a k-nearest neighbors (KNN) graph was constructed using the graph slot from the adjacency matrix of the previously processed *Seurat* object and cells were assigned to neighborhoods (*k=20, d=30*). To leverage the variation in the number of cells between UC, CD and healthy samples, the cells belonging to each sample in each neighborhood were counted and the Euclidean distances between single cells in a neighborhood were calculated. Differential abundance testing in a generalized linear model framework was then performed between IBD samples and healthy controls with default parameters. In addition, to check if the differences in abundances are particularly strong for a certain cell type, we assigned to each neighborhood a specific annotation label considering most cells belonging to that neighborhood. For those neighborhoods where less than 80% of cells shared the most abundant label, a “mixed” label was assigned. Neighborhoods were grouped using the *groupNhoods* function with *max.lfc.delta=5, overlap=3* for CD and healthy samples, and *max.lfc.delta=3, overlap=2* for UC and healthy samples.

***Trajectory analysis***

We applied the Monocle 2 algorithm (v 2.24.1)^15^ to order myeloid cells in pseudotime to indicate their developmental trajectories. Such pseudotime analysis is a measure of progress through biological processes based on their transcriptional similarities. With that aim, we selected all distinct macrophages and monocytes and created a *CellDataSet* object using a negative binomial model. We ran Monocle 2 using the 2000 most highly variable genes selected with *Seurat* (v 4.1.0) and default parameters of Monocle after DDRTree ^16^dimension reduction and cell ordering. To visualize the ordered cells in the trajectory, we used the *plot_cell_trajectory* function to plot the minimum spanning tree on the cells. The starting point of the trajectory was set manually to M0 macrophages.

***Publicly available scRNAseq datasests***

Data from Smillie et al.^17^ and Martin et al.^18^ were downloaded and reanalyzed following the same protocol used for our samples. We separated in silico the macrophage (CD14, CD163, CSF1R, and CD68) compartment to annotate them. In addition to manual annotation, we used the MatchScore2 package^19^ which compares the marker gene list with a reference dataset (in this case our dataset) and assigns a value (Jaccard index) to each comparison. High Jaccard indexes indicate high similarity between clusters. See detailed code [here](https://gitfront.io/r/user-1871750/StAFev2VqZh6/ibd-bcn-single-cell/).

Gene-sets from microarray analysis of stimulated macrophages were obtained from the analysis of GSE121825, GSE155719, GSE156921, GSE161774 and GSE94608 datasets^20-23^. Using these gene-sets, we calculated the average expression levels of each cluster on our dataset at single cell level, using the function *AddModuleScore* of the *Seurat* R Package. See full code [here](https://gitfront.io/r/user-1871750/StAFev2VqZh6/ibd-bcn-single-cell/).

**CosMx™ Spatial Molecular Imaging (SMI)**

CosMx^TM^ technology [CosMx^TM^ Human Universal Cell Characterization RNA Panel (1000-plex) + 20 custom genes; Nanostring, USA] was applied to 9 FFPE samples: 3 non IBD healthy controls, 3 CD and 3 UC with a mean of 19 fields of view (FoVs) per sample. The mean area explored was 11,53 mm^2^ per sample. The following cell surface markers were used for morphology visualization: B2M/CD298, PanCK, CD45, CD3 antibodies and DAPI.

***CosMx™ Spatial Molecular Imager (SMI) sample preparation***

FFPE tissue sections were prepared for CosMx^TM^ SMI profiling as described in He et al^24^. Briefly, five-micron tissue sections mounted on VWR Superfrost Plus Micro slides (cat# 48311-703) were baked overnight at 60°C, followed by preparation for in-situ hybridization (ISH) on the Leica Bond RXm system by deparaffinization and heat-induced epitope retrieval (HIER) at 100°C for 15 minutes using ER2 epitope retrieval buffer (Leica Biosystems product, Tris/EDTA-based, pH 9.0). After HIER, tissue sections were digested with 3 µg/ml Proteinase K diluted in ACD Protease Plus at 40°C for 30 minutes.

Tissue sections were then washed twice with diethyl pyrocarbonate (DEPC)-treated water (DEPC H2O) and incubated in 0.00075% fiducials (Bangs Laboratory) in 2X saline sodium citrate, 0.001% Tween-20 (SSCT solution) for 5 min at room temperature in the dark. Excess fiducials were rinsed from the slides with 1X PBS, then tissue sections were fixed with 10% neutral buffered formalin (NBF) for 5 min at room temperature. Fixed samples were rinsed twice with Tris-glycine buffer (0.1M glycine, 0.1M Tris-base in DEPC H2O) and once with 1X PBS for 5 min each before blocking with 100 mM N-succinimidyl (acetylthio) acetate (NHS-acetate, Thermo Fisher Scientific) in NHS-acetate buffer (0.1M NaP, 0.1% Tween pH 8 in DEPC H2O) for 15 min at room temperature. The sections were then rinsed with 2X saline sodium citrate (SSC) for 5 min and an Adhesive SecureSeal Hybridization Chamber (Grace Bio-Labs) was placed over the tissue.

NanoString® ISH probes were prepared by incubation at 95°C for 2 min and placed on ice, and the ISH probe mix (1nM 980 plex ISH probe, 10nM Attenuation probes, 1nM SMI-0006 custom, 1X Buffer R, 0.1 U/μL SUPERase•In™ [Thermo Fisher Scientific] in DEPC H2O) was pipetted into the hybridization chamber. The chamber was sealed to prevent evaporation, and hybridization was performed at 37°C overnight. Tissue sections were washed twice in 50% formamide (VWR) in 2X SSC at 37°C for 25 min, washed twice with 2X SSC for 2 min at room temperature, and blocked with 100 mM NHS-acetate in the dark for 15 min. In preparation for loading onto the CosMx SMI instrument, a custom-made flow cell was affixed to the slide.

***CosMx Spatial Molecular Imager (SMI) instrument run***

RNA target readout on the CosMx SMI instrument was performed as described in He et al^24^. Briefly, the assembled flow cell was loaded onto the instrument and Reporter Wash Buffer was flowed to remove air bubbles. A preview scan of the entire flow cell was taken, and 15-25 fields of view (FoVs) were placed on the tissue to match regions of interest identified by H&E staining of an adjacent serial section. RNA readout began by flowing 100 μl of Reporter Pool 1 into the flow cell and incubation for 15 min. Reporter Wash Buffer (1 mL) was flowed to wash unbound reporter probes, and Imaging Buffer was added to the flow cell for imaging. Nine Z-stack images (0.8 μm step size) for each FoV were acquired, and photocleavable linkers on the fluorophores of the reporter probes were released by UV illumination and washed with Strip Wash buffer. The fluidic and imaging procedure was repeated for the 16 reporter pools, and the 16 rounds of reporter hybridization-imaging were repeated multiple times to increase RNA detection sensitivity.

After RNA readout, the tissue samples were incubated with a 4-fluorophore-conjugated antibody cocktail against CD298/B2M (488 nm), PanCK (532 nm), CD45 (594 nm), and CD3 (647 nm) proteins and DAPI stain in the CosMx^TM^ SMI instrument for 2 h. After unbound antibodies and DAPI stain were washed with Reporter Wash Buffer, Imaging Buffer was added to the flow cell and nine Z-stack images for the 5 channels (4 antibodies and DAPI) were captured.

***Segmentation and quality control***

Images were segmented to obtain cell boundaries, assign transcripts at the cell-level, and obtain a transcript by cell count matrix^24^. Cells with an average negative control count greater than 0.5 and less than 20 detected features were filtered out. After quality control, a mean of 95,3% of cells across samples and FoVs was retained, corresponding to ~46,160 cells per sample on average.

***Preprocessing and feature selection***

Following library size normalization and log-transformation using *logNormCounts* (*scater* package)^25^, highly variable genes (HVGs) were identified with *modelGeneVar* (*scran* package)^26^, blocking by sample (i.e., variance modelling is performed per sample and statistics combined across samples), and selecting for genes with a positive biological variance component. For comparability with the lowest-coverage sample, per-sample scaling normalization with *multiBatchNorm* from the *batchelor* package was applied^27^.

***Integration and dimension reduction***

Next, Principal Component Analysis (PCA) was computed on highly variable genes (HVGs). Upon inspection of the corresponding elbow plot (% variance explained vs. # of PCs), we selected the first 30 PCs as input for integration using *harmony*^5^, and for dimensionality reduction via uniform manifold approximation and projection (UMAP)^4^

***Cell type annotation***

To identify subpopulation markers, we ran *findMarkers* (*scran* package)^26^ , blocking by sample, considering HVGs only, and testing for positive log-fold changes. The 100 top ranked genes were selected and passed to *SingleR*^28^ along with the reference scRNA-seq dataset’s count matrix and subpopulation assignments, for label transfer. To obtain higher-resolution annotations, we grouped cells into 5 biological compartments according to their lower-resolution label (namely: epithelial, myeloid, plasma, stroma, and T cells) and, for each compartment separately, reperformed marker detection and label transfer as described above.

***Co-localization analysis***

To quantify how pairs of cell types co-localize in a given sample and FoV, we computed, for every sample, FoV, and pair of cell types, two-dimensional kernel density estimates (KDEs) of cell coordinates using *kde*. Within each sample-FoV, estimation was performed over a rectangular window according to the boundary coordinates of cells from a given pair of types, and the Pearson correlation coefficient between KDEs was computed. Only comparisons with more than 10 cells from both types were taken into consideration.

***Data and code availability***

Expression profiles and raw single-cell RNA sequencing data of all cell compartments are publicly available in GEO: [GSE214695](https://www.ncbi.nlm.nih.gov/geo/query/acc.cgi?acc=GSE214695). Analyzed data can be explored in our interactive webpage: <https://servidor2-ciberehd.upc.es/external/garrido/app/>. Full code of the analysis is in our github page <https://github.com/ibd-bcn/ibd-bcn_single_cell> and our main webpage <https://servidor2-ciberehd.upc.es/garrido.html>. CosMx SMI data analysis pipeline available at: <https://github.com/ibd-bcn/CosMx-SMI>. The CosMx SMI data will not be available.

**Granulocytes isolation from blood**

Human granulocytes were isolated from blood by a double density gradient. In brief, diluted blood in PBS (1/2) was layered over Lymphoprep™ (1.077 g/ml) that was layered over denser Polymorphprep™ (1.113 g/ml). The double gradient was then centrifuged at 500g for 30 min obtaining 2 separated cell layers. The lower layer containing the granulocytes was collected, washed and red blood cells were lysed using a commercial lysis buffer (BioLegend). Purity of granulocytes achieved with this method is >95%.

**Flow cytometry**

Blood granulocytes and single-cell suspensions obtained from digested biopsies of heathy donors and IBD patients were stained with the following antibodies CD66b PE-Cy7, CD16 PerCP, CD62L APC-Cy7, CD69 BV421, CD193 FITC, CD63 FITC (BioLegend) and CXCR4 PE (R&D systems) and Zombie Aqua Fixable Viability Kit (BioLegend) for death cells. Cells were fixed using BD Stabilizing Fixative [BD], acquired using a BD FACSCanto II flow cytometer (BD) and analyzed with FlowJO software (BD).

**Organoid cultures from human colonic crypts**

Epithelial 3D organoid cultures were generated using surgical samples from adult healthy donors, as previously described^29^. Ex vivo Organoids embedded in Matrigel (BD Biosciences, CA, USA) were passaged for further expansion approximately every 5-6 days. After 1-3 passages, organoid samples were stimulated with neuregulin 1 (R&D, Germany) at 1 ug/mL for 48h and harvested for RNA extraction.

**RNA isolation and quantitative RT-PCR**

Organoid samples were harvested in Trizol (Ambion, Thermo Fisher Scientific, USA) and RNA was isolated using GeneJET RNA Cleanup and Concentration Micro Kit (Thermo Fisher Scientific, USA). Biopsies were placed in RNAlater RNA Stabilization Reagent (QIAGEN, Hilden, Germany) and stored at –80ºC until RNA isolation. RNA was isolated using an RNeasy Kit (QIAGEN, Hilden, Germany) according to the manufacturer’s instructions and sequenced for bulk RNA-seq. Total RNA from organoid samples were transcribed to cDNA using reverse transcriptase (High Capacity cDNA Archive RT kit, Applied Biosystems, Carlsbad, CA, USA) Quantitative real-time PCR (qPCR) was performed in an ABI PRISM 7500 Fast RT-PCR System (Applied Biosystems) using predesigned TaqMan Assays (Applied Biosystems) (OLFM4, LRG5 and MKI67).

**Bulk RNA-seq**

Barcoded RNAseq libraries were prepared from total RNA using a TruSeq stranded mRNA kit (Illumina, San Diego, CA, USA) according to the manufacturer’s instructions. Libraries were subjected to paired-end sequencing (50 bp) on a HighSeq-3000 platform (Illumina, CA, USA) at the Genomic Service of Boehringer (Boheringer, Germany). Quality filtering and adapter trimming was performed using Skewer version 2.2.8 Reads were mapped against the human reference genome using the STAR aligner version 2.5.2a. The genome used was GRCh38.p10, and gene annotation was based on Gencode version 27 (EMBL-EBI, Hinxton, UK). Read counts per gene were obtained using RSEM version 1.2.31 and the Ensembl GTF annotation file (EMBL-EBI, Hinxton, UK). Analyses were performed using the R (version 3.2.3) statistics package. HC and IBD patient samples used are described in Supplementary Table 1.

**Statistical analysis**

Graphs of bulk RNA-seq, qPCR organoid and flow cytometry results and statistical analysis was performed using Graphpad Prism 9.0 (Graphpad Software, CA, USA). Differences between groups in the bulk RNA-seq data were tested using Ordinary one-way ANOVA test and correcting for multiple comparisons by controlling the false discovery rate [two-stage linear step-up procedure of Benjamini, Krieger and Yekutieli]. Graphs show median. For organoid results, One-sample t test was performed. Data is shown as fold change (FC) relative to the vehicle treated condition. Bars represent mean and standard deviation (SD). Differences between groups in flow cytometry data were tested using the non-parametric Kruskal–Wallis test and correcting for multiple comparisons by controlling the false discovery rate [two-stage linear step-up procedure of Benjamini, Krieger and Yekutieli]. Graphs indicate median and range.

**Immunohistochemistry (IHQ) and immunofluorescence (IHF)**

Immunostaining of tissue sections of healthy donors and IBD patients was performed using commercially available antibodies (anti-rabbit CD68 (Sigma, MO, USA), anti-mouse CD209 (Santa Cruz Biotechnologies, TX, USA), anti-rabbit OLFM4 (Cell Signaling, MA, USA), anti-rabbit MPO (Sigma, MO, USA), anti-mouse MBP (Sigma, MO, USA)). Deparaffinization, rehydration and epitope retrieval of the samples was performed using PT Link (Agilent, CA, USA) using Envision Flex Target Retrieval Solution Low pH (Dako, Germany). Blocking of the samples was performed using horse or goat serum (10-20%) (Vector, NY, USA) in a PBS + 0,5% BSA solution. Secondary antibodies were used for IHQ ( goat anti-rabbit and horse anti-mouse (Vector, NY, USA) and IHF (goat anti-rabbit AF488 and goat anti-mouse Cy3). For IHQ the immunoperoxidase detection system was used (Vector, NY, USA). DAPI (Thermo Fisher Scientific, USA) counterstaining was performed on IHF samples. Image acquisition was performed on a Nikon Ti microscope (Japan) using Nis-Elements Basic Research Software. Immunofluorescence composite was performed using ImageJ software.

**Single molecule RNA *in situ* hybridization (ISH)**

ISH was carried out on FFPE tissues fixed in 10% neutral buffered formalin for 48 hours. RNA probes for NRG1 (cat#311181) and OLFM4 (cat#311041) and RNAscope 2.5 HD assay – Red kit (Biotechne, CO, USA) were used according to the manufacturer’s instructions. Pre-warmed probes were added to the slides and incubated in the HybEZ oven (Biotechne, CO, USA) for 2 hours at 40ºC. After a 6-step signal amplification, for double IHQ and ISH samples, IHQ protocol was followed. Tissue sections were counter-stained with Gill’s hematoxylin. Slides were mounted with ECOmount mounting medium (Biocare medical, CA, USA) and photographed using Nikon Ti microscope (Japan) and Nis-Elements Basic Research Software

<https://pubmed.ncbi.nlm.nih.gov/34594043/>
